# Coordinated hippocampal-thalamic-cortical communication crucial for engram dynamics underneath systems consolidation

**DOI:** 10.1101/2020.12.22.424000

**Authors:** Douglas Feitosa Tomé, Sadra Sadeh, Claudia Clopath

**Affiliations:** Department of Bioengineering, Imperial College London, United Kingdom

## Abstract

Systems consolidation refers to the reorganization of memory over time across brain regions. Despite recent advancements in unravelling engrams and circuits essential for this process, the exact mechanisms behind engram cell dynamics and the role of associated pathways remain poorly understood. Here, we propose a computational model to address this knowledge gap that consists of a multi-region spiking recurrent neural network subject to biologically-plausible synaptic plasticity mechanisms. By coordinating the timescales of synaptic plasticity throughout the network and incorporating a hippocampus-thalamus-cortex circuit, our model is able to couple engram reactivations across these brain regions and thereby reproduce key dynamics of cortical and hippocampal engram cells along with their interdependencies. Decoupling hippocampal-thalamic-cortical activity disrupts engram dynamics and systems consolidation. Our modeling work also yields several testable predictions: engram cells in mediodorsal thalamus are activated in response to partial cues in recent and remote recall and are crucial for systems consolidation; hippocampal and thalamic engram cells are essential for coupling engram reactivations between subcortical and cortical regions; inhibitory engram cells have region-specific dynamics with coupled reactivations; inhibitory input to mediodorsal thalamus is critical for systems consolidation; and thalamocortical synaptic coupling is predictive of cortical engram dynamics and the retrograde amnesia pattern induced by hippocampal damage. Overall, our results suggest that systems consolidation emerges from concerted interactions among engram cells in distributed brain regions enabled by coordinated synaptic plasticity timescales in multisynaptic subcortical-cortical circuits.

## Introduction

Pioneering hippocampal lesion studies [1–3] have motivated an ever-growing body of lesion experiments [4,5] with a common goal of understanding the role of hippocampus and neocortex in systems consolidation of memory. In turn, this spawned many theories of this process but with widely different views concerning its underlying mechanisms and properties. These discrepancies can be mainly attributed to seemingly conflicting reports in the retrograde amnesia literature [4,5]. Specifically, retrograde amnesia induced by hippocampal damage has been reported as temporally-graded (i.e., recent memories are lost but remote memories are spared following hippocampal lesion), flat (i.e., recent and remote memories are disrupted by hippocampal lesion), or absent (i.e., recent and remote memories are preserved post hippocampal lesion). In light of these experimental findings, some systems consolidation theories posited that hippocampus is essential for recent but not for remote memory recall [6–16] while others have proposed that hippocampus is either always necessary for recall [4,17–19] or required for recall depending on the circumstances of encoding and retrieval [20–29]. Despite their differences, these theories share the view that systems consolidation relies on interactions between hippocampus and neocortex. Surprisingly, it has been recently demonstrated that thalamic spindles have a causal role in systems consolidation by coupling hippocampal, thalamic, and cortical oscillations [30]. Therefore, current theories of systems consolidation fail to provide a unifying framework that reconciles the available experimental data.

Recent advances in experimental technologies have the potential to clarify the nature and dynamics of systems consolidation by enabling the identification and manipulation of engrams – more specifically of engram cells [31]. These cells are defined as a set of neurons that become active in response to learning, undergo enduring changes as a result of learning, and are able to be reactivated when presented part of the original stimuli resulting in memory recall [32]. Adopting this definition, a landmark contextual fear conditioning (CFC) study found that engram cells in medial prefrontal cortex (CTX) are initially generated in a silent state (i.e., cannot be reactivated from a partial cue) but over time gradually become active (i.e., can be reactivated from a partial cue) [33]. In contrast, engram cells in hippocampus (HPC) are active following learning but eventually turn silent. The silent-to-active transition in CTX engram cells was named maturation and the active-to-silent switch in HPC engrams was termed de-maturation. Both engram dynamics are associated with systems consolidation of memory. Moreover, the output of HPC engram cells after learning was found to be crucial for the subsequent maturation of CTX engrams. It has been proposed that the observed dynamics of engrams in CTX and HPC [33] are “mirrored” in different types of episodic memory [5]. Nevertheless, the exact neural mechanisms underlying these engram dynamics and the role of associated circuits remain unknown and, consequently, the ability of recent engram findings to advance our knowledge towards a consistent view of systems consolidation is hindered. This is at least in part due to existing theoretical and computational models lagging behind the groundbreaking advancements in engram cell research enabled by new technologies developed in the past decade [31,32,34,35]. In particular, previous computational studies have employed abstract neuronal models that are intended to capture high-level properties of systems consolidation (e.g., recent memory recall relies on hippocampus) but are unable to reproduce engram cell-level data produced by recent experiments [9,11,25,36–40].

Here, our goal is to provide insights into engram cell dynamics and associated pathways using computational modeling. To that end, we simulate systems consolidation in an episodic memory task using a multi-region spiking recurrent neural network model subject to biologically-plausible plasticity mechanisms acting on different timescales in distinct brain regions. Contrary to current theories [4,6–29], our results show that direct, monosynaptic HPC→CTX projections cannot reproduce the known interdependencies between engrams in these regions [33]. However, a network with hippocampal-thalamic-cortical communication is able to overcome this limitation. Specifically, after verifying that our model with three-region communication displays engram cell maturation in CTX and de-maturation in HPC, we then show that HPC engram cells as well as coupled engram reactivations across brain regions are essential for proper engram dynamics in line with previous experiments [30,33]. Our modeling results also yield the following experimentally-testable predictions: engram cells in mediodorsal thalamus (THL) are active in recent and remote recall and are crucial for the maturation of engram cells in CTX; engram cells in HPC and THL are crucial for coupling engram reactivations across HPC, THL, and CTX in consolidation periods; inhibitory engram cells have distinct region-specific dynamics with coupled reactivations; inhibitory input to THL is critical for CTX engram maturation; and THL→CTX synaptic coupling is predictive of CTX engram dynamics and the retrograde amnesia pattern induced by HPC damage — thus providing a unifying mechanistic account for reconciliation of HPC lesion studies. Altogether, our results suggest that coordinated hippocampal-thalamic-cortical communication underlies engram dynamics subserving systems consolidation.

## Results

### Synaptic plasticity timescales drive engram cell dynamics

To understand the mechanisms underlying engram cell dynamics, we start by examining the effects of synaptic plasticity timescales on the initial state and subsequent evolution of engram cells. We use spiking neural network models that consist of a stimulus population (STIM) that projects to both HPC and CTX (Fig. 1**A**). Feedforward and recurrent synapses are initialized at random with excitatory synapses onto excitatory neurons displaying long-term plasticity and inhibitory synapses onto excitatory neurons exhibiting inhibitory plasticity. Long-term excitatory plasticity is composed of a combination of Hebbian and non-Hebbian forms of plasticity [41]. The Hebbian term takes the form of triplet spike-timing-dependent plasticity (STDP) [42] while the non-Hebbian terms include heterosynaptic plasticity [43] and transmitter-induced plasticity [44]. Importantly, the heterosynaptic plasticity term incorporates synaptic consolidation dynamics [41]. Inhibitory synaptic plasticity consists of a network activity-based STDP term [41] whose primary goal is to regulate firing rate levels [45] (for a detailed description of the model, see Methods). One of four non-overlapping random stimuli is presented to the network at a time either for training or testing (Fig. 1**A**), and the network is subject to an episodic memory task to investigate engram dynamics. Following a brief burn-in period to stabilize network activity, the network simulation consists of three consecutive phases: training, consolidation, and testing (Fig. 1**B**). In the training phase, the complete stimuli (i.e., full patterns) are randomly presented to the network. Next, no stimulus is presented to the network during the consolidation phase and, consequently, the network is allowed to evolve spontaneously. At different points throughout the consolidation phase, the network proceeds to the final testing phase where partial cues of the original stimuli are presented and the ability of HPC and CTX to recall the memory is evaluated.

**Figure 1.**
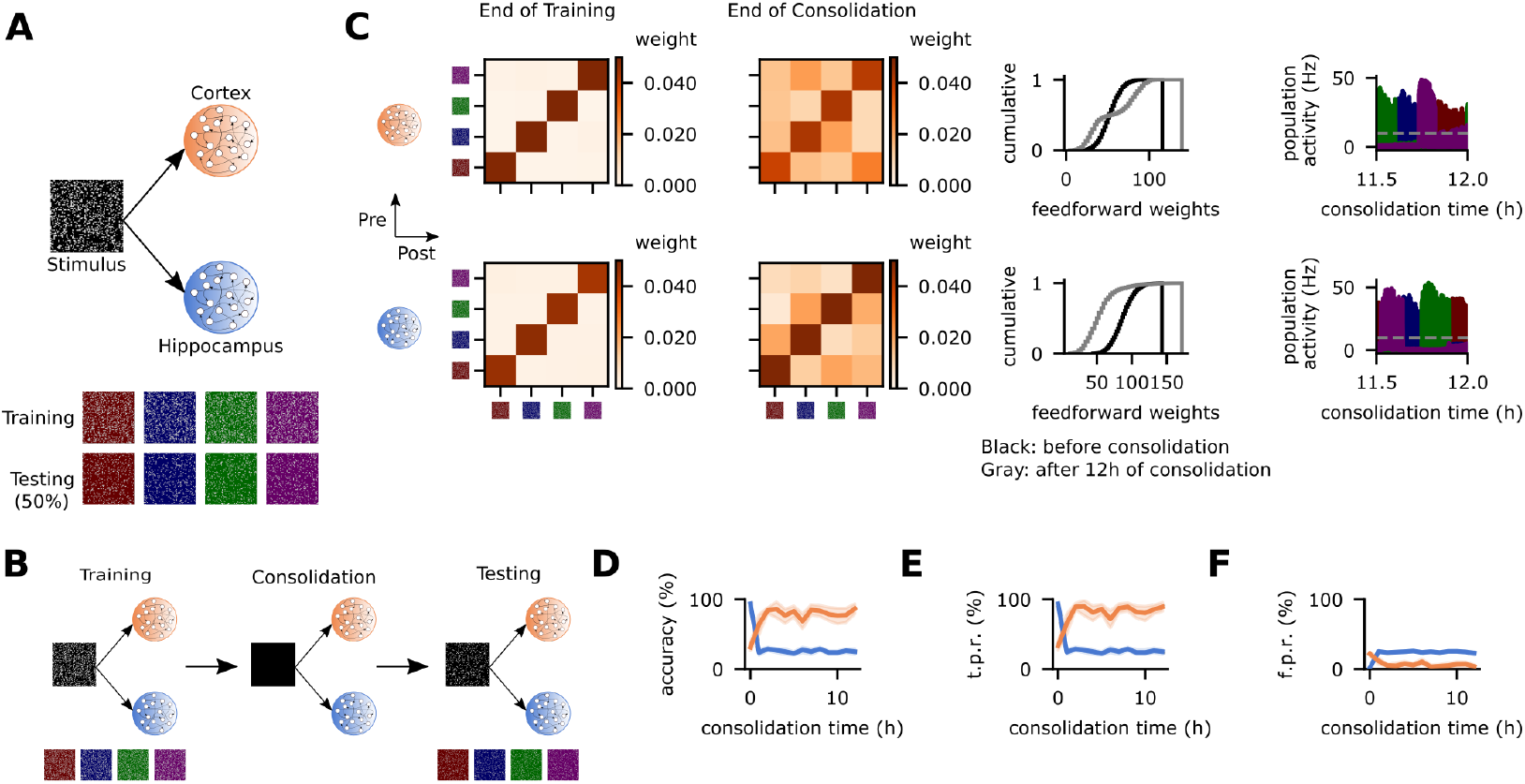
Divergent synaptic plasticity timescales lead to opposite engram cell dynamics. **A**, Schematic of network model with STIM, HPC, and CTX (top) and stimuli presented in the training phase with their respective partial cues used in the testing phase (bottom). **B**, Schematic of simulation protocol. **C**, From left to right: mean weight strength of recurrent excitatory synapses onto excitatory neurons at the end of the training phase clustered according to engram cell preference, mean weight strength of recurrent excitatory synapses onto excitatory neurons after 12 hours of consolidation clustered according to engram cell preference, cumulative distribution function of the total feedforward synaptic weights onto individual engram cells, and population activity of engram cells encoding each stimulus (dashed line indicates threshold *ζ^thr^* = 10 Hz for engram cell activation). Top: CTX. Bottom: HPC. **D-F**, Memory recall in the testing phase of protocol **B** as a function of consolidation time. **D**, Recall accuracy. **E**, Recall true positive rate. **F**, Recall false positive rate. **C-F**, Color as in **A**. **D-F**, *n* = 10 trials with 90% confidence intervals.

Engram cells are formed in both HPC and CTX at the end of training (Fig. 1**C**). These cells are identified via the average stimulus-evoked firing rate of neurons (see Methods). The block diagonal structure (i.e., strong diagonal and weak off-diagonal mean weights) of the recurrent excitatory synapses in HPC and CTX shows that engram cells in these regions have encoded the four stimuli by the end of the training phase. After 12 hours of consolidation, the diagonal structure is preserved in both regions. However, feedforward synapses projecting to HPC and CTX evolve in opposite ways in the consolidation phase. Specifically, while the cumulative distribution function (c.d.f.) of the total feedforward synaptic weights to each engram cell in HPC shows that there is a decrease in STIM→HPC weights, the reverse was observed in CTX. These changes in STIM feedforward weights are consistent with experimental findings which showed that changes in the dendritic spine density of engram cells over time are region-specific with cells in HPC experiencing a decrease but neurons in CTX undergoing an increase in spine density [33]. Furthermore, although engrams are spontaneously reactivated during the consolidation period in the two regions of the model, engram reactivations are not coupled. This was expected given that there are no connections between HPC and CTX (Fig. 1**A**) and, hence, they behave independently in this network configuration. Critically, the differences in engram dynamics in HPC and CTX are a direct result of their diverging synaptic plasticity timescales: learning rate (n) and synaptic consolidation time constant (τ^*cons*^) are higher in HPC relative to CTX.

We next evaluate the ability of the network to retrieve memories from partial cues by plotting memory recall curves. These curves show that the model exhibits de-maturation and maturation of engram cells in HPC and CTX, respectively (Fig **1D-F**), in line with reported experiments [33]. At the end of training (i.e., 0 hours of consolidation), recall true positive rate (t.p.r.) is nearly 100% and false positive rate (f.p.r.) is virtually 0% in HPC, leading to a corresponding recall accuracy of almost 100%. In CTX, however, t.p.r. is approximately 40% and f.p.r. is around 20% with a resulting accuracy of ~40% at the end of encoding. Over the course of the consolidation phase, though, the recall metrics reverse: t.p.r. and accuracy decrease in HPC but they increase in CTX. Importantly, the changes in recall accuracy are driven by changes in t.p.r. in both regions. This means that engram cells in HPC are initially reactivated in response to partial cues but over time become unable to do so while those in CTX cannot be reactivated by cues immediately after training but acquire this ability over the course of consolidation. Consequently, memory recall switches from HPC to CTX with systems consolidation. These engram dynamics are a direct result of region-specific changes in STIM feedforward weights: depression of STIM→HPC synapses and potentiation of STIM→CTX projections (Fig 1**C**). Note that I) HPC engram cells are able to retain their recurrent excitatory structure despite turning silent because of engram reactivations during consolidation (Fig 1**C**) and II) CTX engram cells already have structured recurrent excitatory connectivity at the end of encoding when they are still silent and this enables engram reactivations throughout consolidation (Fig 1**C**). Given the block diagonal structure of the recurrent excitatory weights of silent engram cells in both HPC and CTX, our model suggests that optogenetically stimulating silent engrams in either region triggers memory recall. Interestingly, previous experiments have demonstrated that this is the case [33]. Thus, our modeling results suggest that variability in synaptic plasticity timescales may underlie the observed divergence in engram dynamics across distinct regions of the brain.

### Monosynaptic input from HPC to CTX presents a dilemma

Despite replicating major engram cell dynamics in HPC and CTX, the network model in Fig. 1**A** cannot capture any interdependence between these regions. Given that it has been shown that the output of engram cells in HPC after training is crucial for the maturation of CTX engram cells [33], we now include in our model monosynaptic projections from HPC to CTX with the goal of reproducing this experimental finding. We start by making HPC→CTX synapses plastic with a learning rate 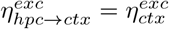 and a synaptic consolidation time constant 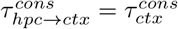 (Fig. 2**A**). This network is also subject to a simulation protocol consisting of training, consolidation, and testing (Fig. 2**B**, compare to Fig. 1**B**). Following training under this configuration, engram cells in HPC and CTX become coupled via the HPC→CTX synaptic weights with a clear diagonal structure (Fig 2**C**). This results in high memory recall accuracy immediately after training in both HPC and CTX (Fig. 2**D**). While engram cells in HPC later undergo de-maturation (similarly to Fig. 1**D**), engram cells in CTX do not display proper maturation. Instead, CTX recall accuracy decreases at the beginning of the consolidation phase and then slowly recovers reaching a level close to the one at the end of training. In order to probe the role of HPC engram cells in the dynamics of CTX engrams, we block the output of HPC engrams during the consolidation period (Fig. 2**E**). In this simulation, the recall accuracy in CTX remains relatively constant with only temporary minor oscillations (Fig. 2**F**). Comparing the simulations with intact and blocked HPC engram cells during consolidation (Fig. **2D** to Fig. 2**F**), it is clear that, in this configuration with plastic HPC→CTX synapses (Fig. 2**A**), engram cells in HPC do not support CTX engram maturation and are effectively detrimental to remote recall in CTX. This is due to input from HPC acting as noise after its engram cells have de-matured. Thus, this network configuration (Fig. 2**A**) neither captures engram cell maturation in CTX nor the crucial role that engram cells in HPC play in this process.

**Figure 2.**
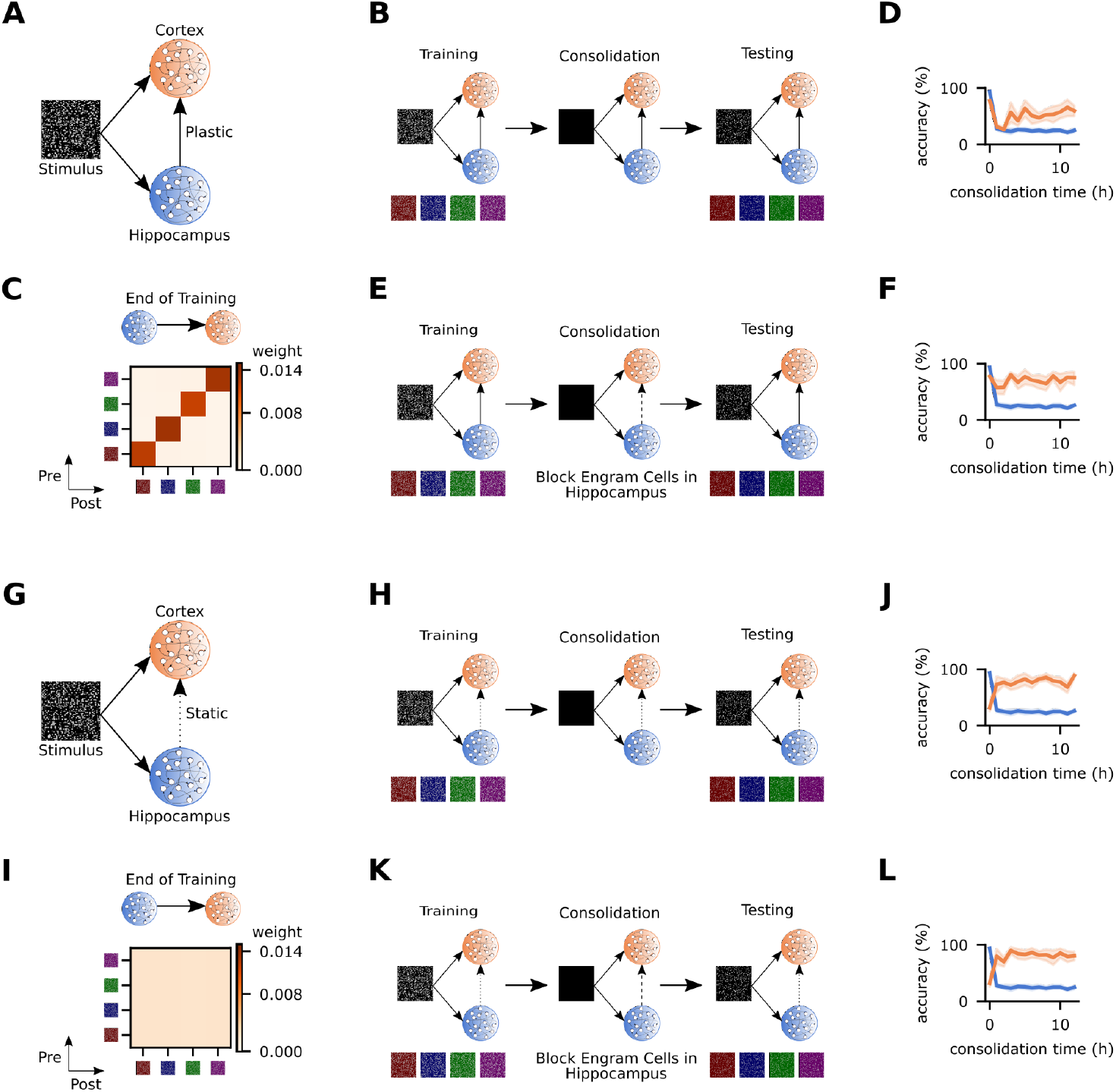
Direct HPC→CTX coupling has contradictory effect. **A/G**, Schematic of network model where HPC→CTX synapses are plastic (**A**) and static (**G**). **B/H**, Schematic of simulation protocol with intact (control) HPC→CTX synapses for networks **A** and **G**, respectively. **C/I**, Mean HPC→CTX weight strength at the end of training clustered according to engram cell preference for networks **A** and **G**, respectively. **D/J**, Memory recall accuracy in the testing phase of protocols **B** and **H**, respectively. **E/K**, Schematic of simulation protocol with the output of engram cells in HPC blocked during consolidation for networks **A** and **G**, respectively. **F/L**, Memory recall accuracy in the testing phase of protocols **E** and **K**, respectively. **D/F**, Color as in **A**. **J/L**, Color as in **G**. **B-C/E/H-I/K**, Stimuli as in Fig. 1 **A**. **D/F/J/L**, *n* = 10 trials with 90% confidence intervals.

Alternatively, we consider a network where the synapses from HPC to CTX are static (Fig. 2**G**). Similarly to the protocol applied to the configuration with plastic HPC→CTX synapses (Fig. 2**B**), this network undergoes training, consolidation, and testing (Fig. 2**H**). In this configuration, the mean weight matrix of HPC→CTX synapses at the end of training shows that differences in the mean weight strength between engrams in these two regions are random and virtually nonexistent and, hence, there is no functional synaptic coupling between them (Fig. 2**I**). This network exhibits proper engram dynamics (i.e., de-maturation in HPC and maturation in CTX) as shown by its recall accuracy curves with intact HPC→CTX synapses (Fig. 2**J**). However, blocking the output of engram cells in HPC during consolidation (Fig. 2**K**) reveals that they are not necessary for the maturation of CTX engrams in this network: CTX recall accuracy with blocking (Fig. 2**L**) has the same trend of silent to active engram cells as without blocking (Fig. 2**J**). This result was predictable since the lack of synaptic plasticity between HPC and CTX and the consequent absence of functional synaptic coupling between engrams in these two regions make the input from HPC to CTX act as a source of noise that does not bear any active role in the dynamics of CTX engram cells. Therefore, static HPC→CTX synapses cannot reproduce the interdependence between these two regions despite accurately capturing their specific engram cell state transitions. The network configurations with static HPC→CTX synapses (Fig. 2**G**) and without HPC→CTX projections (Fig. 1**A**) have essentially the same behavior (compare Fig. **2J** to Fig. 1**D**). Importantly, we explored a wide range of alternative synaptic plasticity timescales (e.g., Fig. **S1A-C**) and network configurations including STIM, HPC, and CTX (e.g., Fig. **S1D-J**) and found that they consistently yield networks with HPC→CTX synaptic coupling that varies between the two extremes discussed previously: strong coupling (Fig. 2**C**) and absent coupling (Fig. 2**I**). These intermediate levels of HPC→CTX synaptic coupling result in HPC engram cells having an impact on CTX engram dynamics that oscillates between strongly detrimental (Fig. **2D/F**) and negligible (Fig. **2J/L**). Taken together, these results leave us with a dilemma: neither plastic nor static monosynaptic projections from HPC to CTX can capture both the engram cell dynamics and interdependencies seen in experiments [33].

### Subcortical engram cells are essential for CTX engram cell maturation

To find a solution to the dilemma previously presented (Fig. 2), we re-examined the brain regions that provide monosynaptic input to engram cells in CTX. Specifically, it has been reported that in CFC the ventral hippocampus (vHPC) is the only hippocampal area that has direct projections to CTX engram cells, but this amounts to only ~5% of their total monosynaptic input [33]. This led us to hypothesize that HPC engram cells use a multisynaptic pathway to CTX to support the maturation of its engrams. In order to test this hypothesis, we include THL in our model since it simultaneously I) receives input from HPC (via the medial temporal lobes: entorhinal and perirhinal cortices [46,47]), II) has a large share of the monosynaptic projections to CTX engram cells (~20%) [33], III) is essential for remote memory recall in CFC [48], and IV) has increased activity around hippocampal ripples coupled to spindles [49] – noting that the latter have a causal role in the systems consolidation of an episodic memory [30]. As a result, we expand the network with THL and set plastic and static circular receptive fields in STIM→HPC and STIM→THL, respectively (Fig. 3**A**). We then use a different set of stimuli for training (i.e., four non-overlapping horizontal bars) and testing (i.e., the central 50% of each bar) (Fig. 3**A**). In this network configuration, HPC and CTX are readout populations but not THL. This means that memory recall from a partial cue is only considered successful if it can be retrieved in either HPC or CTX. We then set the learning rates 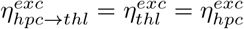, and 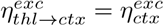 with 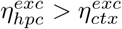); reflecting that subcortical synapses tend to change at a faster rate than their cortical counterparts. In line with our previous results (Fig. 1), STIM→HPC synapses have a longer synaptic consolidation time constant 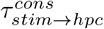 relative to the other excitatory projections in the network (for further details, see Methods). Altogether, this network configuration allows us to evaluate whether the HPC→THL→CTX multisynaptic circuit can provide a pathway for HPC engram cells to support the maturation of CTX engrams.

**Figure 3.**
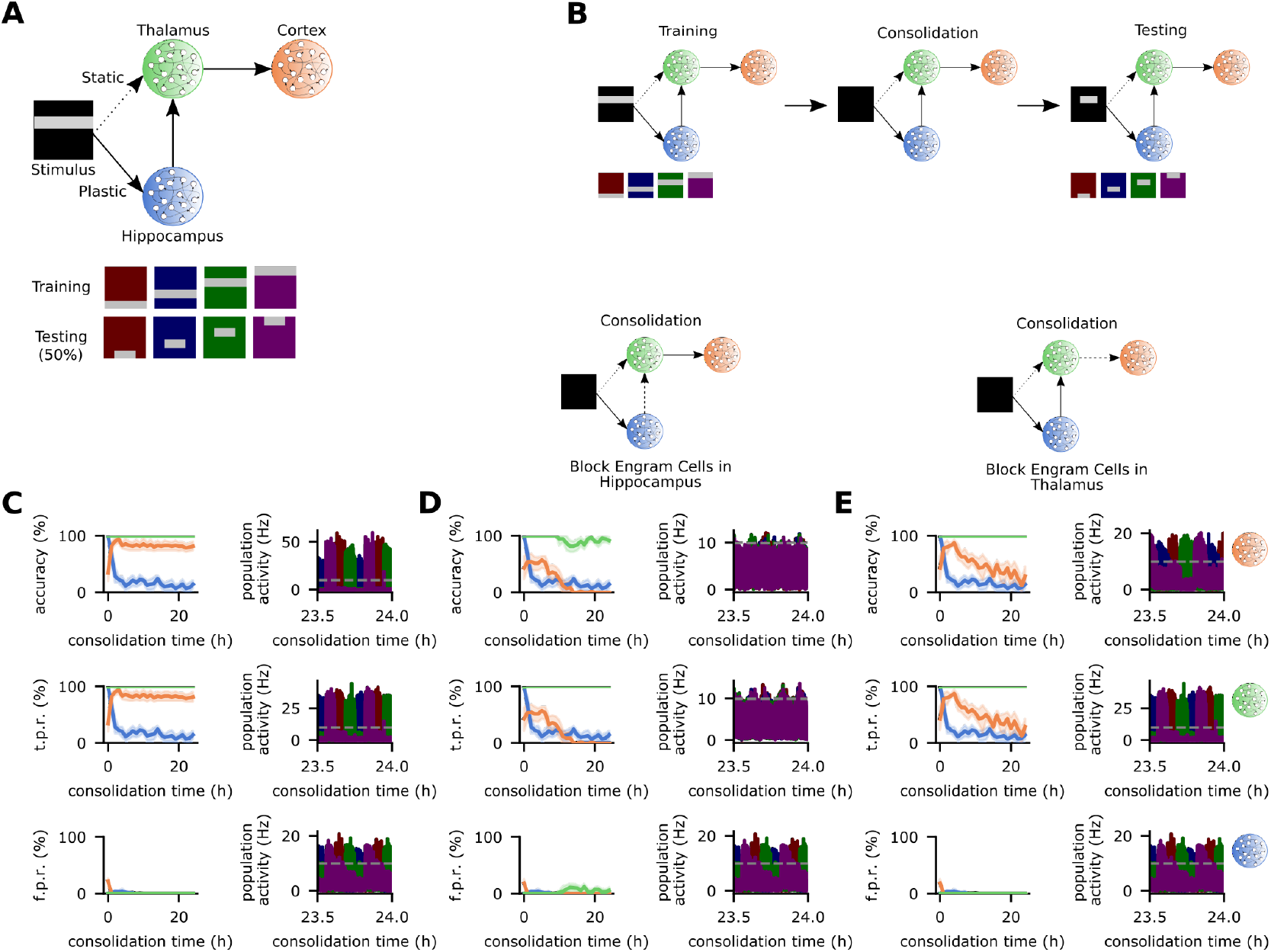
Engram cells in HPC and THL are crucial for the maturation of CTX engram cells. **A**, Schematic of network model with HPC, THL, and CTX (top) and stimuli presented in the training phase with their respective partial cues used in the testing phase (bottom). STIM→THL synapses are static but the remaining feedforward projections are plastic. **B**, Schematic of simulation protocol. **C-E**, Memory recall in the testing phase as a function of consolidation time (left) and population activity in the consolidation phase (right). Recall curves (top to bottom): accuracy, true positive rate, and false positive rate. Population activity of engram cells (top to bottom): CTX, THL, and HPC (dashed line indicates threshold *ζ^thr^* = 10 Hz for engram cell activation). **C**, intact (control) network as in **B**. **D**, output of engram cells in HPC blocked during consolidation. **E**, output of engram cells in THL blocked during consolidation. **B-E**, Color as in **A**. **C-E**, Left: *n* = 5 trials with 90% confidence intervals.

We then subject the three-region network (Fig. 3**A**) to training, consolidation, and testing (Fig. 3**B**) and verify that it also exhibits de-maturation and maturation of engram cells in HPC and CTX, respectively (Fig. 3**C**). Hence, memory recall switches from HPC to CTX with consolidation (Fig. S2 and S3) due to changes in engram cell state that are driven by changes in t.p.r. This is consistent with previous findings [33] and reflects region-specific plastic changes in feedforward afferent synapses: depression of STIM→HPC projections and potentiation of THL→CTX synapses (Fig. S4) in a manner analogous to the two-region network (Fig. 1**C**). Engram cells in THL are initially active and remain so throughout the consolidation period in our simulations. Importantly, excitatory and inhibitory plasticity are required for proper engram dynamics (Fig. S5). Furthermore, the engram dynamics observed in HPC, THL, and CTX are accompanied by coupled engram reactivations across these three regions (Fig. 3**C**). Precise coupling of oscillations in HPC, THL, and CTX in consolidation periods has been previously shown to be linked to systems consolidation [30,50–54]. Therefore, our model exhibits engram dynamics and oscillation patterns consistent with previous experiments.

We next probe the role of HPC engram cells in the maturation of CTX engrams. To that end, we block the output of engram cells in HPC during consolidation and subsequently test memory recall (Fig. 3**D**). Although recall accuracy in CTX initially shows a modest increase, it goes on to suffer a sharp decline and eventually settles at nearly zero. This is driven by the CTX t.p.r. curve which displays the same pattern. Hence, engram cells in HPC are crucial for the maturation of CTX engram cells in the three-region network and reflect previous findings [33]. Plotting the population activity in the network reveals that the blockage of HPC engram cells disrupts the coupling of engram reactivations in the consolidation phase (Fig. 3**D**). Recent experiments that tampered with the coupling of oscillations in HPC, THL, and CTX have demonstrated that this has an adverse effect on the systems consolidation of episodic memory [30, 50]. Thus, our model suggests that engram cells in HPC are essential for the maturation of CTX engrams because they support coupling engram reactivations across brain regions throughout consolidation periods.

We then evaluate whether THL engram cells are also critical for CTX engram maturation. In a manner analogous to our previous probe of HPC engram cells, we block the output of engram cells in THL in the consolidation phase. The resulting memory recall curves show that THL engram cells are also crucial for the maturation of engram cells in CTX (Fig. 3**E**). The CTX recall accuracy curve in this case has a different shape though: at first accuracy increases substantially and reaches a level similar to the control network (Fig. 3**C**) before gradually declining and finally reaching a low point after more than 20 hours of consolidation (compare this to Fig. 3**D**). This trend in recall accuracy is driven by CTX t.p.r. and suggests that THL engram cells are essential for stabilizing active engram cells in CTX. Despite the differences in recall profile between blocking HPC and THL engram cells, both lead to uncoupled engram reactivations between CTX and the remaining subcortical regions in the network (Fig. **3D-E**). As expected, blocking the output of THL engram cells does not affect the coupling of reactivations in HPC and THL. Nevertheless, this has a downstream effect on the THL-CTX coupling leading to decoupled reactivations in these regions. Taken together, our modeling results predict that THL engram cells are essential for CTX engram cell maturation as they also support the coupling of subcortical-cortical oscillations in a manner similar to HPC engram cells.

We can gain a deeper understanding of the mechanisms underlying the active role of HPC and THL engram cells in the maturation of CTX engrams by examining changes in synaptic weights in the network. At the end of training, engrams in HPC, THL, and CTX are coupled via the feedforward synapses in the HPC→THL→CTX circuit (Fig. 4**A**). THL and CTX recurrent excitatory weights form a block diagonal structure that reflects stimulus encoding. After 24 hours of consolidation, HPC→THL coupling is degraded but a clear separation between on- and off-diagonal weights still persists while THL→CTX connectivity is not only preserved but reinforced (Fig. 4**B**). The recurrent weight structures of THL and CTX are also largely maintained. Despite de-maturation, HPC engram cells also retain a learning-induced block diagonal structure after consolidation (Figure S6) similarly to the two-region network (Fig. 1**C**). In contrast, blocking HPC engram cells during consolidation greatly damages the diagonal pattern in both feedforward and recurrent weights (compare Fig. **4C** and **B**). HPC→THL coupling and THL recurrent weights are the most affected with nearly no distinction between on- and off-diagonal elements but the impact on THL→CTX coupling is also significant. This results in decoupled oscillations in HPC, THL, and CTX (Fig. 3**D**). This effect is compounded by the degradation of the recurrent excitatory weight structure in CTX (Fig. 4**C**) and the subsequent deterioration of engram reactivations in this region (Fig. 3**D**). Although blocking THL engram cells has no impact on the evolution of HPC→THL and recurrent THL weights as expected (compare Fig. **4D** and **B**), it compromises THL→CTX coupling in a similar way as did blocking HPC engram cells (compare Fig. **4D** and **C**). Accordingly, this also leads to decoupled reactivations between THL and CTX (Fig. 3**E**). A deteriorating effect on CTX recurrent weights is also visible (Fig. 4**D**) but in this case it does not prevent reactivations in CTX altogether (compare Fig. **3E** and **D**). Thus, engram cells in HPC and THL are crucial for maintaining strong synaptic coupling in the HPC→THL→CTX pathway and, consequently, coordinating engram reactivations across these regions to support CTX engram maturation.

**Figure 4.**
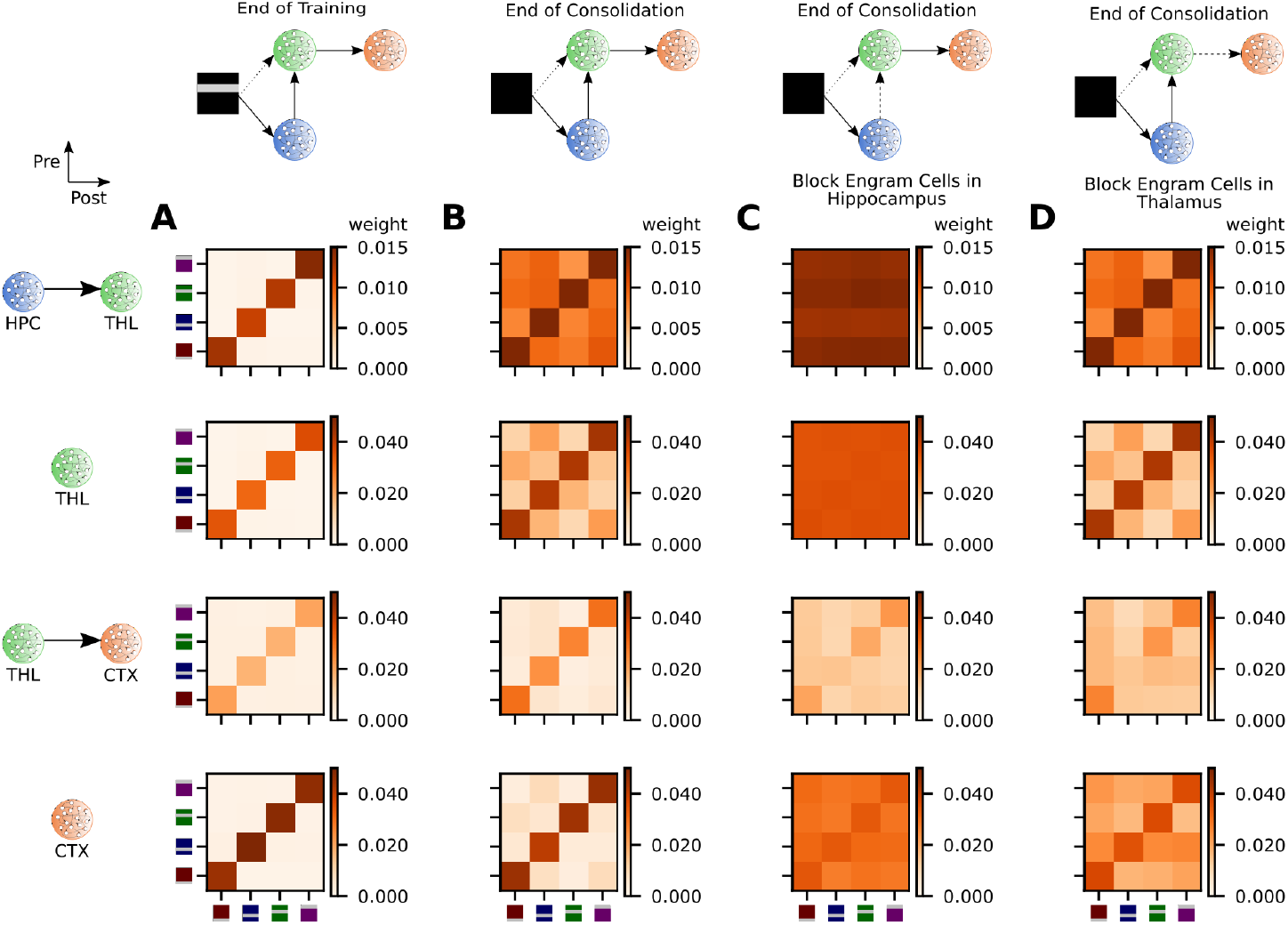
Subcortical engram cells are essential for the consolidation of thalamocortical coupling. **A-D**, Mean weight strength of excitatory synapses onto excitatory neurons clustered according to engram cell preference for the network in Fig. 3**A**. From top to bottom: HPC→THL (feedforward), THL (recurrent), THL→CTX (feedforward), and CTX (recurrent). **A**, Mean weight matrices at the end of the training phase (Fig. **3C-E**). **B**, Mean weight matrices after 24 hours of consolidation for the intact (control) network (Fig. 3**C**). **C**, Mean weight matrices after 24 hours of consolidation for the network with blocked HPC engram cells (Fig. 3**D**). **D**, Mean weight matrices after 24 hours of consolidation for the network with blocked THL engram cells (Fig. 3**E**).

### Inhibitory engram cells have distinct dynamics

Engram cell experiments have focused on excitatory neurons given that expression of immediate early genes (IEGs) used for activity-dependent labelling occurs predominantly in these cells [35,55,56]. However, growing evidence suggests that inhibitory engrams co-exist with excitatory engram cells [57,58]. We therefore also investigate the behavior of inhibitory neurons in our model. We start by comparing the recall profile of inhibitory and excitatory engram cells (Fig. **5A** and Fig. 3**C**, respectively). The CTX recall accuracy of both sets of engram cells increases over the consolidation period but this rise is driven by a decrease in f.p.r. in the case of inhibitory neurons while it is caused by an increase in t.p.r. for excitatory cells. Effectively, inhibitory engram cells have a high t.p.r. post-training with only minor subsequent oscillations but excitatory engrams have a flat near-zero f.p.r. throughout consolidation. Therefore, inhibitory engram cells in CTX become stimulus-specific with consolidation whereas excitatory engrams become active. The sharpening of the response of CTX inhibitory engrams can be attributed to potentiation of their inhibitory synapses onto excitatory engram cells in the consolidation period (Fig. 5**B**). In addition, the recall accuracy of THL inhibitory engram cells immediately after training is at 100% and then quickly decays to zero due to a sharp uptake in f.p.r. while t.p.r. remains at 100%. This means that THL inhibitory engram cells are continuously active after encoding but become progressively more unspecific to stimuli as a result of depression of their inhibitory synapses projecting to excitatory engrams in the consolidation phase (Fig. 5**B**). This is in stark contrast to THL excitatory engrams which remain active and stimulus-specific after training. Furthermore, excitatory and inhibitory engram cells in HPC undergo de-maturation in a cascading manner: excitatory engram cells coding a given stimulus become silent in the consolidation period and, consequently, the corresponding inhibitory engrams are no longer activated by partial cues either. Note that de-maturation of HPC inhibitory engrams is accompanied by potentiation of their inhibitory synapses onto excitatory engram cells (Fig. 5**B**). Lastly, excitatory and inhibitory engram cells have different composition profiles (Fig. S7). Overall, the dynamics of inhibitory engram cells in our model vary by brain region.

**Figure 5.**
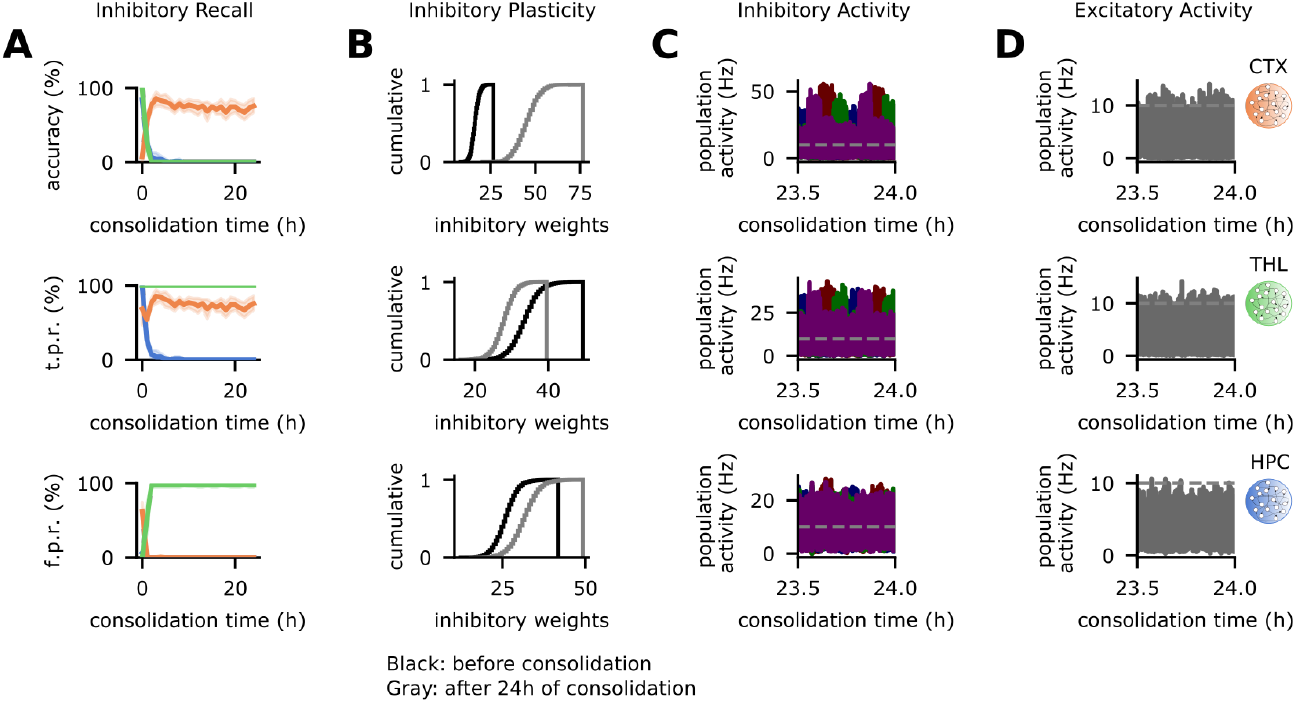
Dynamics of inhibitory engram cells are region-specific. **A-D**, Analysis of engram dynamics in protocol shown in Fig. 3**B**. **A**, Recall of inhibitory engram cells in the testing phase as a function of consolidation time. From top to bottom: accuracy, true positive rate, and false positive rate. **B**, Cumulative distribution function of the total inhibitory synaptic weights onto individual excitatory engram cells at the end of training and after 24 hours of consolidation. **C-D**, Population activity in the consolidation phase (dashed line indicates threshold *ζ^thr^* = 10 Hz for engram cell activation). **C**, Population activity of inhibitory engram cells. **D**, Population activity of excitatory neurons. **B-D**, From top to bottom: CTX, THL, and HPC. **A/C-D**, Color as in Fig. 3**A**. **A**, *n* = 5 trials with 90% confidence intervals.

Inhibitory engram cells in HPC, THL, and CTX also have coupled reactivations in the consolidation phase (Fig. 5**C**, S8). Note that the activity of inhibitory engrams in the three regions remains coupled despite the fact that the amplitude of HPC inhibitory oscillations becomes progressively smaller as consolidation progresses. Comparing the activity of inhibitory and excitatory engram cells (Fig. **5C** and Fig. 3**C**, respectively), we can see that reactivations are coordinated across neuron types in each region throughout consolidation (Fig. S8). The oscillatory activity of inhibitory engram cells combined with inhibitory synaptic plasticity (Fig. 5**B**) is able to tame the activity of excitatory neurons in each individual area (Fig. 5**D**, S9) while still allowing coupled reactivations of excitatory engram cells. Taken together, our results predict that inhibitory engram cells have region-specific dynamics and that reactivations of inhibitory and excitatory engrams are coupled in consolidation periods.

### Inhibitory input to HPC, CTX, and THL is crucial for CTX engram dynamics

We probe the role of region-specific inhibitory input in engram dynamics throughout the network by blocking the output of inhibitory neurons during consolidation. Specifically, we first block the output of HPC inhibitory neurons in the consolidation phase and this disrupts CTX engram maturation and the coupling of engram reactivations in the HPC→THL→CTX circuit (Fig. 6**A**). Blocking inhibitory neurons in CTX also tampers with CTX engram dynamics and the subcortical-cortical coupling of engram reactivations (Fig. 6**B**). These results are aligned with previous findings that showed that blocking parvalbumin-positive interneurons either in HPC or CTX decoupled oscillations in these regions in consolidation periods and disrupted systems consolidation [50]. We then block inhibitory neurons in THL and this also prevents engram cell maturation in CTX and the coupling of engram reactivations in the network (Fig. 6**C**). In each of the previous simulations, blocking inhibitory input to one region significantly alters the dynamics of engrams in that region and in any downstream areas (Fig. **6A-C**) as inhibitory drive is essential for the consolidation of subcortical-cortical synaptic coupling (Fig. S10). Our model then predicts that inhibitory input to HPC, CTX, and THL is essential for CTX engram maturation by coupling engram reactivations in consolidation periods. Given that blocking excitatory engram cells in HPC and THL (Fig. **3D-E**) and blocking inhibitory input to HPC, CTX, and THL (Fig. **6A-C**) both decouple engram reactivations across these regions and disrupt CTX engram maturation, our results suggest that coordinated HPC-THL-CTX communication underlies engram dynamics that mediate systems consolidation.

**Figure 6.**
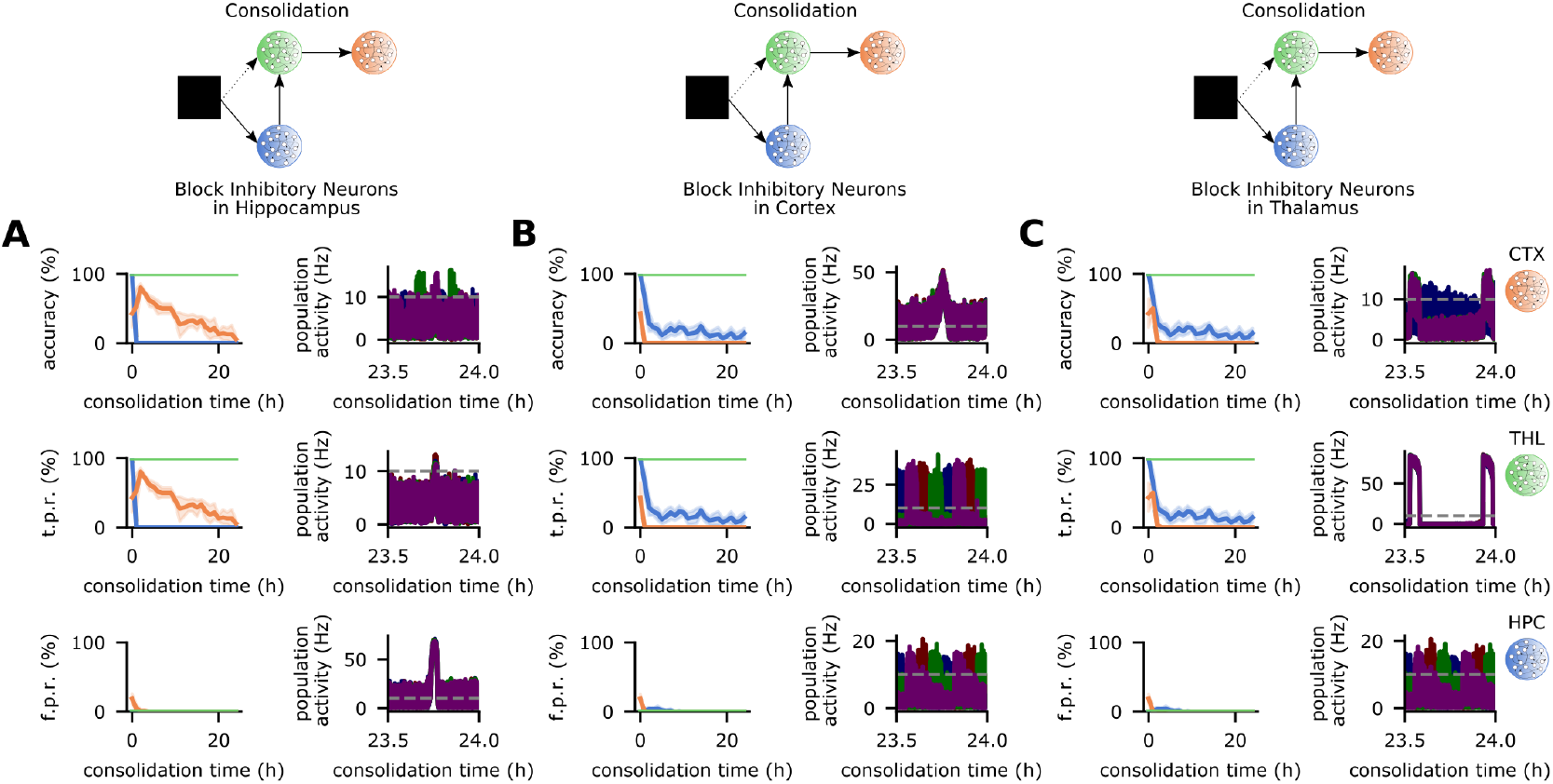
Inhibitory input to HPC, CTX, and THL is critical for CTX engram maturation. **A-C**, Memory recall as a function of consolidation time (left) and population activity in the consolidation phase (right) when region-specific inhibitory neurons are blocked in the consolidation phase of the protocol depicted in Fig. 3**B**. Recall curves (top to bottom): accuracy, true positive rate, and false positive rate. Population activity of excitatory engram cells (top to bottom): CTX, THL, and HPC (dashed line indicates threshold *ζ^thr^* = 10 Hz for engram cell activation). **A**, output of inhibitory neurons in HPC blocked during consolidation. **B**, output of inhibitory neurons in CTX blocked during consolidation. **C**, output of inhibitory neurons in THL blocked during consolidation. **A-C**, Color as in Fig. 3**A**. **A-C**, Left: *n* =5 trials with 90% confidence intervals.

### Thalamocortical coupling underlies retrograde amnesia profiles

We then investigate to what extent memory recall relies on HPC over time by examining retrograde amnesia patterns induced by HPC ablation. Ablation of HPC in the testing phase (Fig. 7**A**) leads to significant impairment in recent recall (Fig. 7**B**) since it originally relied on HPC (Fig. 3**C**). However, remote recall is virtually not affected by HPC ablation (Fig. 7**B**). Therefore, memory recall reliance on HPC is time-dependent and the model exhibits a temporally-graded retrograde amnesia curve [4].

**Figure 7.**
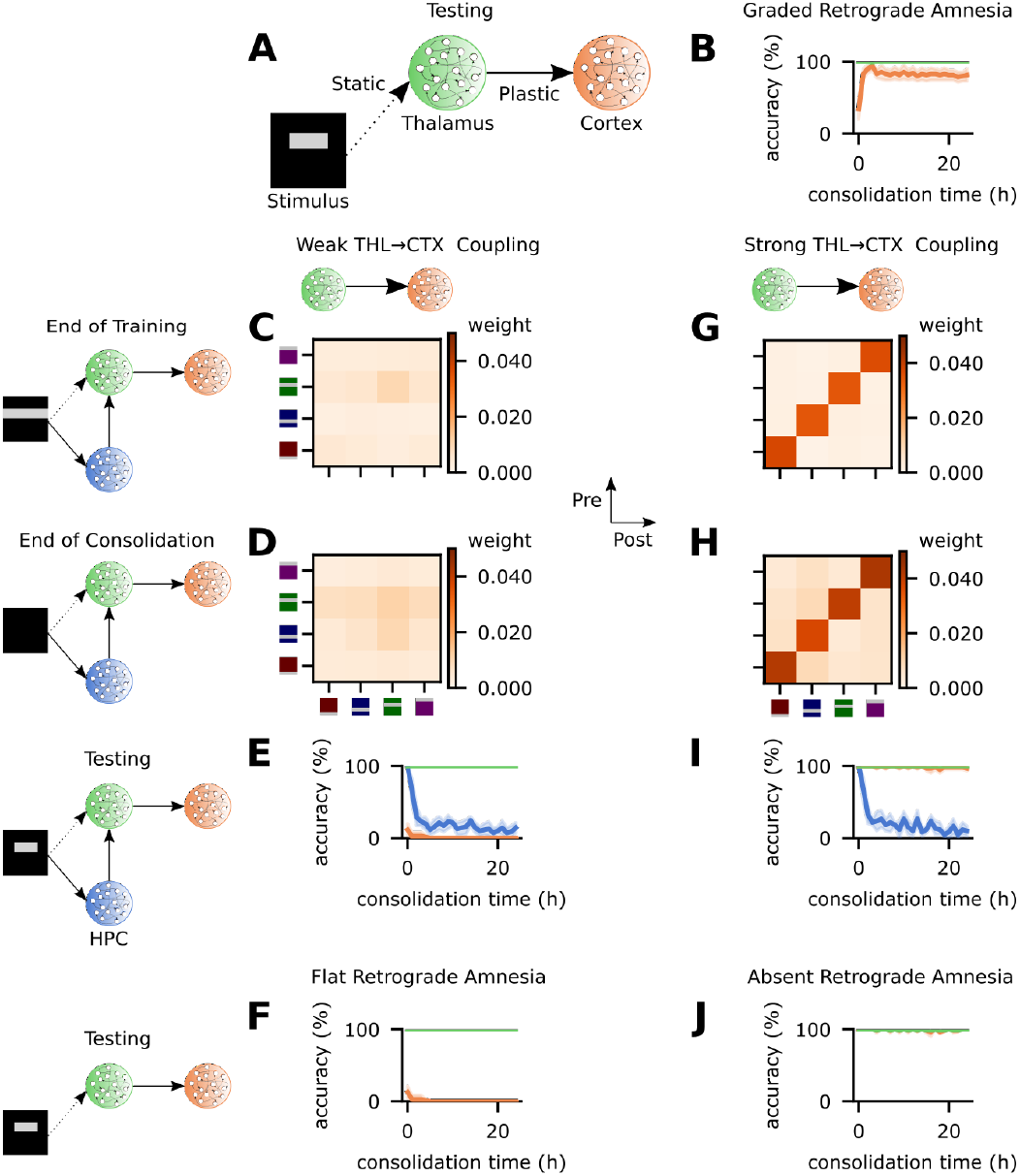
THL→CTX coupling at the end of encoding underlies retrograde amnesia pattern. **A**, Schematic of network model with ablation of HPC at testing time. **B**, Memory recall accuracy as a function of consolidation time with HPC ablation in the testing phase (**A**) of protocol shown in Fig. 3**B**. **C-F**, Simulation with uncoupled THL→CTX. **G-J**, Simulation with strongly coupled THL→CTX. **C-D/G-H**, Mean weight strength of THL→CTX synapses clustered according to engram cell preference. **C/G**, At the end of training. **D/H**, After 24 hours of consolidation. **E/I**, Memory recall accuracy as a function of consolidation time with intact network in protocol depicted in Fig. 3**B**. **F/J**, Memory recall accuracy as a function of consolidation time with HPC ablation in the testing phase (**A**) of protocol shown in Fig. 3**B**. **B-J**, Color as in Fig. 3**A**. **B/E-F/I-J**, *n* = 5 trials with 90% confidence intervals.

We next probe the role of THL→CTX coupling on HPC reliance by varying the plasticity rate of these synapses. Specifically, we explored how heterosynaptic plasticity strength *β_THL→CTX_* can increase or decrease coupling between THL and CTX at the end of encoding and the resulting effect on memory recall. We increase *β_THL→CTX_* substantially (Fig. **7C-F**), which, consequently, severely impairs the ability of THL→CTX synapses to potentiate (see Methods). As a result, THL and CTX are uncoupled at the end of the training phase (Fig. 7**C**, compare to Fig. 4**A**) and remain so despite subsequent consolidation (Fig. 7**D**). Accordingly, remote recall with the intact control network (Fig. 3**A**) is lost since the decoupling of THL→CTX synapses prevents CTX engram maturation and HPC engram cells still become silent (Fig. 7**E**). Naturally, HPC ablation does not improve remote recall and it also prevents recent memory retrieval (Fig. 7**F**). Thus, the network with uncoupled THL→CTX synapses relies exclusively on HPC for memory recall and displays a flat retrograde amnesia pattern [4].

Subsequently, we reduce *β_THL→CTX_* to effectively enable faster synaptic potentiation (see Methods) and, consequently, increase THL→CTX synaptic coupling at the end of encoding (Fig. 7**G**, compare to Fig. 4**A**). Coupling between these regions is reinforced with consolidation (Fig. 7**H**) and CTX recall accuracy is therefore extremely high both immediately following training and throughout consolidation (Fig. 7**I**). Ablating HPC has a negligible effect on CTX recall and, hence, memories can be recalled independently of HPC (Fig. 7**J**). This network configuration then exhibits an absent retrograde amnesia curve [4]. Altogether, our model predicts that the degree of THL→CTX synaptic coupling at the end of encoding is a major driver of the ensuing CTX engram cell dynamics and associated retrograde amnesia profiles induced by HPC ablation.

## Discussion

Our model is able to reproduce key experimental findings associated with systems consolidation. Specifically, it captures engram cell maturation and de-maturation in CTX and HPC, respectively, and the crucial role that HPC engram cells have in the maturation of CTX engrams [33]. The model also reflects the causal role of coupled oscillations across HPC, THL, and CTX in the systems consolidation of episodic memory [30] and connects it to the associated engram dynamics observed in experiments [33]. We have demonstrated that these experimental findings can be reproduced in a computational model of the HPC→THL→CTX multisynaptic pathway with region-specific synaptic plasticity rates. Our results suggest that the timescale of synaptic plasticity is precisely conducted across brain regions to enable coordinated HPC-THL-CTX communication and that these concerted subcortical-cortical interactions are vital for engram dynamics behind systems consolidation of memory.

The timescales of the various forms of synaptic plasticity in our model need to be coordinated to reproduce specific engram cell state transitions taking place in parallel. The learning rate of the triplet STDP is higher in subcortical regions (i.e., HPC and THL) relative to CTX consistent with the view that subcortical synapses tend to be more plastic than cortical ones. However, synaptic consolidation is slower in HPC compared to THL and CTX in line with the observation that HPC engram cells are less stable and, hence, more prone to becoming silent. Interestingly, it has been previously suggested that synaptic consolidation has an active role in systems consolidation [59]. Transmitter-induced plasticity rates are scaled linearly to the learning rate of each individual region to prevent long-term depression (LTD) from making the network silent while the timescales of heterosynaptic plasticity are set to avoid excessive network activity while still allowing long-term potentiation (LTP) to take place. This combination of Hebbian (triplet STDP) and non-Hebbian (heterosynaptic and transmitter-induced) plasticity has been shown to enable stable memory formation and recall in a single-region spiking neural network model [41] and here we build on those results to show that coordinated synaptic plasticity timescales across brain regions can extend the mnemonic functions supported by these forms of plasticity.

There are multiple circuits that can potentially be used by HPC to support the maturation of CTX engram cells but we include the HPC→THL→CTX pathway in our model. As noted earlier, this choice is motivated by the afferent and efferent projections of THL (i.e., HPC and CTX, respectively [46,47,60]) and the observation that in CFC this region is both responsible for ~20% of the monosynaptic input to CTX engram cells [33] and crucial for remote recall [48]. Here, we assume that THL also has an essential role in the remote recall of other types of episodic memories in a similar way as it has been proposed that engram cell dynamics observed in CFC are present in generic episodic memories [5] — a view that is consistent with numerous reports of memory impairments in a wide range of tasks following lesions to THL [46,47]. Furthermore, the increased THL activity around hippocampal ripples coupled to spindles [49] suggests that this region may play a part in the essential role that spindles have in coupling cortical, thalamic, and hippocampal oscillations in systems consolidation [30]. Thus, the HPC→THL→CTX circuit seems to be a prime candidate for having a crucial role in the maturation of CTX engram cells and our modeling results support this view. Nevertheless, we cannot exclude the possibility that alternative circuits may also be used by HPC for the same purpose. In fact, three other brain regions have monosynaptic projections to CTX engram cells to a similar extent as THL in CFC: anterodorsal thalamus (ADT), medial entorhinal cortex layer Va (MEC-Va), and basolateral amygdala (BLA) [33]. Note, however, that I) ADT is only essential for recent but not for remote CFC memory recall [48], II) MEC-Va→CTX is not required for neither recent nor remote recall in CFC [33], and III) BLA→MEC stimulation improved retention of the contextual but not foot shock components of memory in CFC [61]. In addition, a multisynaptic pathway involving the dorsoventral axis of HPC may also be used by HPC engram cells to support engram dynamics in CTX given that the dorsal hippocampus (dHPC) has a critical role in CFC [62,63]. Hence, dHPC→vHPC→CTX may be recruited by HPC engrams but as noted earlier in CFC only ~5% of the total monosynaptic input to CTX engram cells originates in vHPC [33]. Further, although another possible circuit may involve dHPC and restrosplenial cortex (RSC) (i.e., dHPC→RSC→CTX), RSC projections only account for less 10% of the monosynaptic input to CTX engram cells in CFC [33]. Altogether, these findings pose HPC→THL→CTX as a plausible minimal circuit for the encoding, consolidation, and recall of episodic memory and our simulation results support this viewpoint.

The mechanisms through which HPC engram cells support the maturation of CTX engrams remain unknown but our modeling results suggest that HPC engrams are essential for coupling hippocampal, thalamic, and cortical engram reactivations and thereby are crucial for CTX engram cell maturation. The causal role of coupled HPC-THL-CTX oscillations in the systems consolidation of episodic memories has been previously demonstrated [30, 50] and our model predicts that HPC engram cells have themselves a causal role in this coupling. Although engram cells have been found in various thalamic nuclei [64], the potential role that thalamic engram cells have in systems consolidation is not known either. Our simulations suggest that engram cells in THL are also crucial for the maturation of CTX engram cells by coupling engram reactivations across HPC, THL, and CTX. Importantly, two lines of evidence support the view that engrams are present in THL: I) a large body of THL lesion studies that showed post-lesion memory deficits in a diverse array of tasks [46, 47, 65]; and II) a recent experiment that found THL to be one of the regions with a high probability of holding an engram in a brain-wide mapping of CFC memory [64]. Furthermore, our model predicts that THL engram cells are active in both recent and remote recall in a similar way as BLA engram cells in CFC [33].

Our model also aims to shed light on the dynamics of inhibitory engram cells. “Inhibitory replicas” of learning-induced excitatory connectivity patterns have been found [58, 66], but the behavior of inhibitory engram cells has not been probed yet. Our model predicts that the dynamics of inhibitory engrams are region-specific: CTX inhibitory engram cells are active in recent and remote recall but become more selective to stimuli over time, THL inhibitory engrams also maintain an active status after training but rather become unspecific to stimuli, and HPC inhibitory engram cells undergo de-maturation similarly to their excitatory counterparts. Inhibitory engrams are formed in our model via the potentiation of inhibitory synapses onto excitatory cells that display learning-induced activity increase in line with other computational models [57, 66]. Furthermore, our results also predict reactivations of inhibitory engram cells coupled to oscillations of excitatory engrams in HPC, THL, and CTX. Previously, inhibitory neurons were shown to control the size of excitatory engram cell ensembles [67, 68] and to mediate memory discrimination [69]. Here, we suggest interneurons also undergo learning-induced changes akin to excitatory neurons (i.e., inhibitory engrams). Moreover, blocking inhibitory neurons in HPC and CTX disrupts systems consolidation in our simulations by preventing coordinated communication between these regions. This is consistent with the crucial role of inhibitory neurons in coupling CTX spindles and HPC ripples [50]. Our model also predicts that inhibitory input to THL has a similar critical role in coupling engram reactivations across subcortical and cortical regions. Importantly, local THL inhibitory neurons are present in primates but not in lower species such as rodents [47]. However, the thalamic reticular nucleus (TRN) provides robust inhibitory input to THL across species via GABAergic projections [47, 70]. For those species that rely exclusively on TRN for inhibitory control of network activity, TRN inhibitory neurons may play an analogous role to that of local THL interneurons in higher species given that I) TRN was shown to have a high probability of holding engram cells in a brain-wide mapping of CFC in rodents [64] and II) TRN has an active role in the generation of thalamocortical oscillatory rhythms [70, 71]. Altogether, our results suggest that inhibitory neurons in distributed brain regions have a crucial role in the coordination of HPC-THL-CTX communication mediated by engram reactivations.

We also investigate how recent and remote recall rely on HPC by reproducing the different patterns of retrograde amnesia induced by HPC damage: temporally-graded, flat, and absent [4]. Our model predicts that the degree of THL→CTX synaptic coupling at the end of encoding is predictive of the subsequent CTX engram cell dynamics (i.e., silent or active at recent and remote recall) and the corresponding retrograde amnesia profile caused by HPC lesions. Our model then also predicts that silent CTX engram cells are the basis of retrograde amnesia induced by HPC damage. This is consistent with protein synthesis inhibitor-induced retrograde amnesia studies that showed that silent HPC engram cells underlie this form of amnesia and that their afferent synapses from upstream engram cells exhibit reduced potentiation relative to active engram cells in healthy mice [72, 73]. Furthermore, the discovery of a rapidly-encoded engram in human posterior parietal cortex [74] suggests the existence of cortical engram cells that are active in recent and remote recall as predicted by our model. Taken together, our modeling results predict that distinct engram cell dynamics underlie specific patterns of retrograde amnesia induced by HPC damage and, thus, provide a mechanistic account to reconcile seemingly conflicting reports in the HPC lesion literature [4, 5].

Our model makes several testable predictions. First, our results predict that engram cells in THL are active in recent and remote recall and are crucial for the maturation of engram cells in CTX. This could be tested by labelling THL engram cells during encoding and subsequently blocking their output in a manner analogous to previous protocols [33]. Second, our model predicts that engram cells in HPC and THL are essential for coupling engram reactivations in HPC, THL, and CTX in consolidation periods. Blocking separately HPC and THL engram cells after training [33] and measuring the degree of coupling of oscillations in HPC-THL-CTX during sleep [30] could test this prediction. Third, our results suggest that inhibitory engram cells with region-specific dynamics and coupled reactivations co-exist with excitatory engrams in subcortical and cortical regions. Although engram cell studies have focused on excitatory neurons due to their increased IEG expression [35, 55, 56] as discussed previously, inhibitory neurons can also up-regulate IEGs (e.g., c-fos and Arc) under strong stimulation [35,75,76]. Therefore, our predictions regarding inhibitory engram cells could also be tested by modifying parameters of existing engram cells experiments [33] to induce reliable IEG expression in inhibitory neurons and, hence, enable labelling of inhibitory engram cells. Fourth, our model predicts that inhibitory input to THL is critical for CTX engram maturation by coupling engram reactivations in subcortical-cortical circuits. This prediction could be tested by extending current engram cell protocols [33] with chemogenetic techniques already used to block interneurons in HPC and CTX [50] but applying them to inhibitory neurons projecting to THL. Fifth, our model suggests that synaptic coupling in THL→CTX at the end of encoding is predictive of CTX engram dynamics and the resulting pattern of retrograde amnesia induced by HPC damage. These predictions could also be tested by incorporating activity-dependent cell labelling – combined with strategies for circuit-specific manipulations and in vivo calcium imaging – to existing HPC lesion protocols. THL→CTX synaptic coupling could potentially be either decreased by applying protein synthesis inhibitors to CTX [72,73] or increased by extending total training time and/or stimulus exposure.

Despite capturing engram cell dynamics in HPC and CTX and the coupling of oscillations in the HPC→THL→CTX circuit, our model has several limitations. First, systems consolidation takes place over days, months, or even years after memory acquisition [5] but our simulations extend for only 24 hours after training. Although the synaptic plasticity rates in our model could conceivably be reduced to match more realistic timescales of systems consolidation, this would immensely increase the computational cost of simulations and, consequently, it would be impractical to simulate multi-region large-scale networks like ours for long periods. Second, we do not explicitly model hippocampal sharp-wave ripples, thalamic spindles, and cortical slow oscillations. Instead, our model captures the fact that oscillations in HPC, THL, and CTX need to be coupled for effective systems consolidation [30] by displaying coupled engram cell reactivations. Third, we do not attempt to model a gradual shift in memory from episodic (i.e., specific, detail-rich) to semantic (i.e., abstract, gist-like) over systems consolidation. The extent of such change is still an open question [5] and is beyond the scope of the present study.

In the long history of the field, many computational models of systems consolidation have been proposed [9,11,36–40]. While early computational studies relied on networks with highly abstract, simplified neuron models [9,11,36,37], recent computational models have become increasingly more complex to incorporate a wider range of experimental findings: a three-stage Bayesian Confidence Propagation Neural Network was used to bridge the gap between working and long-term memory [38], a spiking network was developed to explore the role of anatomical properties of the cortex-hippocampus loop in systems consolidation [39], and a rate-coded multi-layer network with a form of Hebbian learning was employed to investigate the effect of preexisting knowledge on memory consolidation [40]. Nevertheless, previous models have not addressed recent findings regarding engram cells and their role in systems consolidation. Our work, however, reproduces engram cell dynamics in HPC and CTX in a novel computational model – specifically, a multi-region spiking recurrent neural network with biologically-plausible synaptic plasticity. In addition, our model also reflects the role of coupled oscillations across HPC, THL, and CTX in systems consolidation and connects it to engram cell reactivations.

In conclusion, our model of systems consolidation exhibits known region-specific engram cell dynamics and captures the active role of both HPC engram cells and coupled HPC-THL-CTX oscillations in this process. We also make several testable predictions regarding HPC and THL engram cells, inhibitory engram cells, inhibitory input to THL, and the relationship between THL→CTX synaptic coupling and retrograde amnesia induced by HPC lesions. Overall, our results suggest that coordinated communication across subcortical-cortical circuits — enabled by coupled engram reactivations — is essential for engram dynamics that ultimately culminate in systems consolidation. Engram cell dynamics in other brain regions, engram interactions in multi-task settings, and the link between engram cells and neurodegenerative diseases will each warrant future experimental and computational studies.

## Methods

### Neuron model

Our model makes use of leaky integrate-and-fire neurons with spike frequency adaption. The membrane voltage *U_i_* of neuron *i* evolves according to [41]:

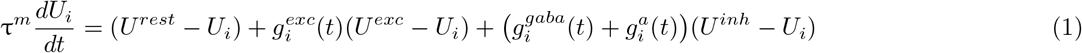

where *τ^m^* is the membrane time constant, *U^rest^* is the membrane resting potential, *U^exc^* is the excitatory reversal potential, and *U^inh^* is the inhibitory reversal potential. The evolution of the synaptic conductance terms 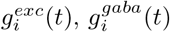, and 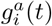 is discussed in the next section.

A neuron *i* fires a spike when its membrane voltage exceeds a threshold *ϑ_i_*. At this point, its membrane voltage is set to 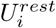 and its firing threshold is temporarily increased to *ϑ^spike^*. Without further spikes, the firing threshold decays to its resting value *ϗ^rest^* with time constant *τ^thr^* following:

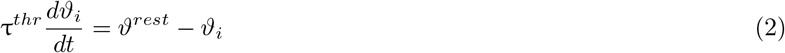

### Synapse model

We adopted a conductance-based synaptic input model. The dynamics of inhibitory synaptic input 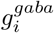 and spike-triggered adaption 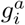 follow [41]:

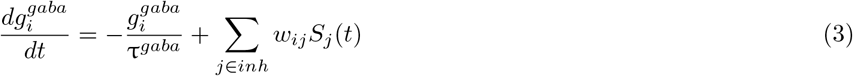

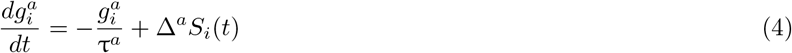

where 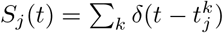 is the presynaptic spike train and 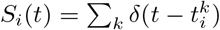 is the postsynaptic spike train. In both cases, *δ* denotes the Dirac delta function and 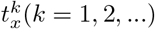 are the firing times of neuron *x. w_ij_* is the weight from neuron *j* to neuron *i*. Δ^*a*^ is a fixed adaptation strength. *τ^gaba^* is the GABA decay time constant and τ^*a*^ is the adaptation time constant.

Excitatory synaptic input is determined by a combination of a fast AMPA-like conductance 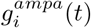 and a slow NMDA-like conductance 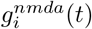:

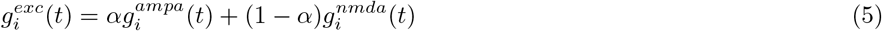

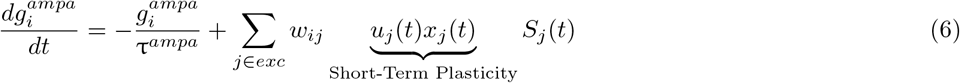

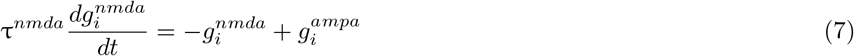

where *α* is a constant that determines the relative contribution of 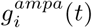 and 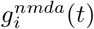 while *τ^ampa^* and *τ^nmda^* are their respective time constants. *u_j_* (*t*) and *x_j_*(*t*) are variables that determine the state of short-term plasticity as described in the following section.

### Synaptic plasticity model

Our synaptic plasticity model was designed after previous work that showed that a combination of Hebbian (i.e., triplet STDP) and non-Hebbian (i.e., heterosynaptic and transmitted-induced) forms of plasticity can yield stable memory formation and recall in a single-region spiking recurrent neural network [41].

#### Short-term plasticity

The state variables *u_j_*(*t*) and *x_j_*(*t*) associated with short-term plasticity evolve according to [41]:

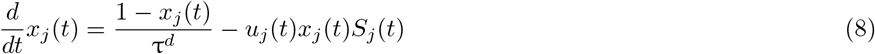

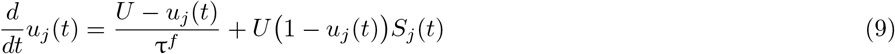

where *τ^d^* and *τ^f^* are the depression and facilitation time constants, respectively. The parameter *U* is the initial release probability.

#### Long-term excitatory synaptic plasticity

Long-term excitatory synaptic plasticity takes the form of combined triplet STDP [42], heterosynaptic plasticity [43], and transmitter-induced plasticity [44] with a synaptic weight *w_ij_* from neuron *j* to neuron *i* following [41]:

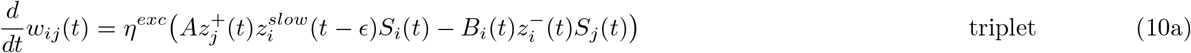

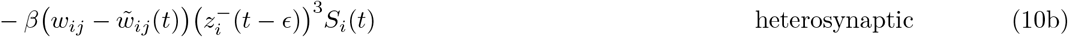

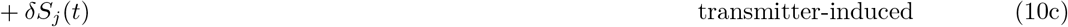

where *η^exc^* (excitatory learning rate), *A* (LTP rate), *β* (heterosynaptic plasticity strength), and *δ* (transmitter-induced plasticity strength) are fixed parameters. *ϵ* is an infinitesimal offset used to ensure that the current action potential is not considered in the trace. State variables 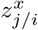 denote either pre- or postsynaptic traces and each has an independent temporal evolution with time constant *τ^x^* given by:

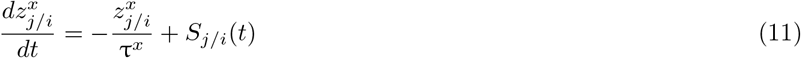

The reference weights 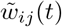 also have their own independent synaptic consolidation dynamics [41]:

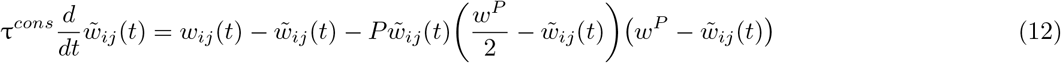

where *P* and *w^P^* are fixed parameters and *τ^cons^* is a time constant. Importantly, the LTD rate *B_i_*(*t*) is subject to homeostatic regulation and evolves according to:

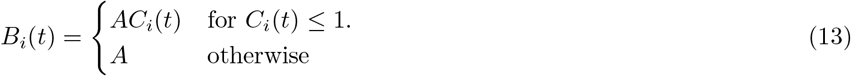

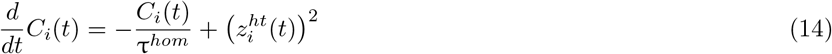

where *τ^hom^* is a time constant and 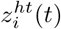 is a synaptic trace that follows Equation 11 with its own time constant *τ^ht^*. Lastly, plastic excitatory weights are constrained to lower and upper bounds 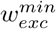 and 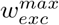, respectively. However, excitatory weights never reach their upper bound with the exception of some simulations with blockage of neurons.

#### Inhibitory synaptic plasticity

Inhibitory synaptic plasticity follows a network activity-based STDP rule [41]:

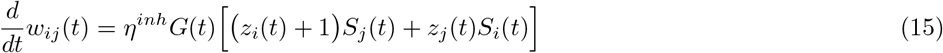

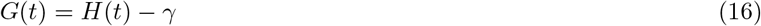

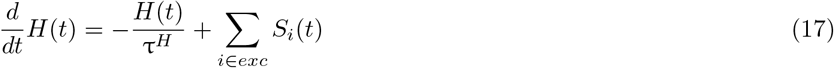

where *η^inh^* is a constant inhibitory learning rate and *z_j/i_* denotes either pre- or postsynpatic traces that follow Equation 11 with a common time constant *τ^iSTDP^*. *G*(*t*) is a linear function of the difference between a hypothetical global secreted factor *H*(*t*) and the target local network activity level *γ*. *H*(*t*) is itself a low-pass-filtered version of the spikes fired by all excitatory neurons in the local network (i.e., either HPC, THL, or CTX) with time constant *τ^H^*. Note that inhibitory synaptic plasticity primarily aims to control network activity. Finally, inhibitory weights are constrained to the interval between 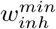 and 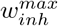 but they never reach their upper limit except in some simulations with blockage of neurons.

### Network model

In each network configuration considered (i.e., Fig. 1**A**, Fig. **2A/G**, and Fig. 3**A**), the model consists of a stimulus population of *N_stim_* = 4, 096 Poisson neurons (STIM) and two or three subnetworks each corresponding to a different brain region (i.e., HPC, CTX, or THL). Each region subnetwork is composed of *N_exc_* = 4, 096 excitatory neurons and *N_inh_* = 1,024 inhibitory neurons that are recurrently connected. Recurrent excitatory synapses onto excitatory neurons display short- and long-term excitatory synaptic plasticity while excitatory synapses projecting onto inhibitory neurons exhibit only short-term plasticity. Feedforward inter-region synapses may display both short- and long-term plasticity or only short-term plasticity depending on the network configuration and they project exclusively from excitatory neurons in one region to excitatory cells in another area. In the two-region networks (Fig. 1**A**, Fig. **2A/G**), all recurrent and feedforward synapses are initialized at random following a uniform distribution. In the three-region network (Fig. 3**A**), STIM→HPC and STIM→THL synapses have randomly-centered circular receptive fields (i.e., each excitatory neuron in HPC and THL receives projections from a small circular area in STIM of radius *R_hpc_* and *R_thl_*, respectively, whose random center location follows a uniform distribution) but the remaining feedforward as well as all recurrent synapses are initially random following a uniform distribution. In addition, inhibitory synapses onto inhibitory neurons are static while inhibitory synapses projecting onto excitatory neurons display inhibitory synaptic plasticity. Plasticity is constantly active for the entirety of all simulations. Recurrent synapses are connected with probability *ϵ_rec_* and are initialized with specific weights (i.e., *w^EE^*, *w^EI^*, *w^II^*, and *w^IE^*). Feedforward synapses have specific connection probabilities and initial weights (e.g., *ϵ_hpc→ctx_* and *w_hpc→ctx_,* respectively, for Fig.**2A/G**). For a complete list of network parameters, see Table 1.

**Table 1.** Summary of network simulation parameters.

| Neural Populations |  |  |  |
| --- | --- | --- | --- |
| Parameter | Figure | Value | Description |
| $N_{exc}$ | All | 4096 | Size of population of excitatory neurons in cortex, hippocampus, and thalamus |
| $N_{inh}$ | All | 1024 | Size of population of inhibitory neurons in cortex, hippocampus, and thalamus |
| $N_{stim}$ | All | 4096 | Size of population of stimulus neurons |
| $N_{ext}^{ctx}$ | 3,4,5,6,7 | 4096 | Size of population of external background neurons projecting to cortex |
| $N_{ext}^{hpc}$ | 3,4,5,6,7 | 4096 | Size of population of external background neurons projecting to hippocampus |

| Network Connectivity |  |  |  |
| --- | --- | --- | --- |
| Parameter | Figure | Value | Description |
| $\epsilon_{rec}$ | All | 0.05 | Probability of connection of recurrent synapses (EE, EI, II, IE) |
| $\epsilon_{stim}$ | 1,2 | 0.1 | Probability of connection of STIM→CTX and STIM→HPC synapses |
| $\epsilon_{hpc \rightarrow ctx}$ | 2 | 0.02 | Probability of connection of HPC→CTX synapses |
| $R_{hpc}$ | 3,4,5,6,7 | 8 | Radius of receptive field of excitatory neurons in hippocampus (STIM→HPC) |
| $R_{thl}$ | 3,4,5,6,7 | 4 | Radius of receptive field of excitatory neurons in thalamus (STIM→THL) |
| $\epsilon_{hpc \rightarrow thl}$ | 3,4,5,6,7 | 0.02 | Probability of connection of HPC→THL synapses |
| $\epsilon_{thl \rightarrow ctx}$ | 3,4,5,6,7 | 0.05 | Probability of connection of THL→CTX synapses |
| $\epsilon_{ext \rightarrow ctx}$ | 3,4,5,6,7 | 0.1 | Probability of connection of CTX external synapses ($EXT_{ctx} \rightarrow CTX$) |
| $\epsilon_{ext \rightarrow hpc}$ | 3,4,5,6,7 | 0.1 | Probability of connection of HPC external synapses ($EXT_{hpc} \rightarrow HPC$) |
| $w^{EE}$ | All | 0.1 | Initial weight of recurrent excitatory synapses onto excitatory neurons |
| $w^{EI}$ | All | 0.6 | Fixed weight of recurrent excitatory synapses onto inhibitory neurons |
| $w^{II}$ | All | 0.2 | Fixed weight of recurrent inhibitory synapses onto inhibitory neurons |
| $w^{IE}$ | All | 0.2 | Initial weight of recurrent inhibitory synapses onto excitatory neurons |
| $w_{stim}$ | 1,2 | 0.1 | Initial weight of STIM→CTX and STIM→HPC synapses |
| $w_{hpc \rightarrow ctx}$ | 2 | 0.1 | Initial weight of HPC→CTX synapses |
| $w_{stim \rightarrow hpc}$ | 3,4,5,6,7 | 0.5 | Initial weight of STIM→HPC synapses |
| $w_{stim \rightarrow thl}$ | 3,4,5,6,7 | 2.0 | Fixed weight of STIM→THL synapses |
| $w_{hpc \rightarrow thl}$ | 3,4,5,6,7 | 0.1 | Initial weight of HPC→THL synapses |
| $w_{thl \rightarrow ctx}$ | 3,4,5,6,7 | 0.15 | Initial weight of THL→CTX synapses |
| $w_{ext \rightarrow ctx}$ | 3,4,5,6,7 | 0.2 | Fixed weight of CTX external synapses ($EXT_{ctx} \rightarrow CTX$) |
| $w_{ext \rightarrow hpc}$ | 3,4,5,6,7 | 0.2 | Fixed weight of HPC external synapses ($EXT_{hpc} \rightarrow HPC$) |

| Neuron Model |  |  |  |
| --- | --- | --- | --- |
| Parameter | Figure | Value | Description |
| $\tau^m$ | All | 20 ms | Membrane time constant |
| $U^{rest}$ | All | -70 mV | Membrane resting potential |
| $U^{exc}$ | All | 0 mV | Excitatory reversal potential |
| $U^{inh}$ | All | -80 mV | Inhibitory reversal potential |
| $\tau^{thr}$ | All | 5 ms | Threshold time constant |
| $\vartheta^{rest}$ | All | -50 mV | Threshold resting value |
| $\vartheta^{spike}$ | All | 100 mV | Threshold value immediately after spike |

| Synapse Model |  |  |  |
| --- | --- | --- | --- |
| Parameter | Figure | Value | Description |
| $\tau^{gaba}$ | All | 10 ms | GABA decay time constant |
| $\tau^a$ | All | 100 ms | Adaptation time constant |
| $\Delta^a$ | All | 0.1 | Adaptation strength |
| $\alpha^E$ | All | 0.2 | AMPA/NMDA ratio for excitatory neurons |
| $\alpha^I$ | All | 0.3 | AMPA/NMDA ratio for inhibitory neurons |
| $\tau^{ampa}$ | All | 5 ms | AMPA decay time constant |
| $\tau^{nmda}$ | All | 100 ms | NMDA decay time constant |

| Short-Term Plasticity Model |  |  |  |
| --- | --- | --- | --- |
| Parameter | Figure | Value | Description |
| $\tau_{EE}^d$ | All | 150 ms | Depression time constant for excitatory synapses onto excitatory neurons |
| $\tau_{EI}^d$ | All | 200 ms | Depression time constant for excitatory synapses onto inhibitory neurons |
| $\tau^f$ | All | 600 ms | Facilitation time constant for excitatory synapses |
| $U$ | All | 0.2 | Initial release probability for excitatory synapses |

| Long-Term Excitatory Synaptic Plasticity Model |  |  |  |
| --- | --- | --- | --- |
| CTX EE |  |  |  |
| Parameter | Figure | Value | Description |
| $\eta_{ctx}^{exc}$ | All | $1 \times 10^{-3}$ | Learning rate of EE synapses in cortex |
| $\lambda_{ctx}^{\beta}$ | All | 50 | $\beta_{ctx}/\eta_{ctx}^{exc}$ ratio for EE synapses in cortex |
| $\tau_{ctx}^{cons}$ | All | 20 min | Synaptic consolidation time constant for EE synapses in cortex |
| HPC EE |  |  |  |
| Parameter | Figure | Value | Description |
| $\eta_{hpc}^{exc}$ | 1,2 | $1.25 \times 10^{-3}$ | Learning rate of EE synapses in hippocampus |
| | 3,4,5,6,7 | $1.5 \times 10^{-3}$ | |
| $\lambda_{hpc}^{\beta}$ | 1,2 | 25 | $\beta_{hpc}/\eta_{hpc}^{exc}$ ratio for EE synapses in hippocampus |
|  | 3,4,5,6,7 | 50 |  |
| $\tau_{hpc}^{cons}$ | 1,2 | 3 h | Synaptic consolidation time constant for EE synapses in hippocampus |
|  | 3,4,5,6,7 | 20 min |  |
| STIM→CTX |  |  |  |
| Parameter | Figure | Value | Description |
| $\eta_{stim \rightarrow ctx}^{exc}$ | 1,2 | $1 \times 10^{-3}$ | Learning rate of STIM→CTX synapses |
| $\lambda_{stim \rightarrow ctx}^{\beta}$ | 1,2 | 85 | $\beta_{stim \rightarrow ctx}/\eta_{stim \rightarrow ctx}^{exc}$ ratio for STIM→CTX synapses |
| $\tau_{stim \rightarrow ctx}^{cons}$ | 1,2 | 20 min | Synaptic consolidation time constant for STIM→CTX synapses |
| STIM→HPC |  |  |  |
| Parameter | Figure | Value | Description |
| $\eta_{stim \rightarrow hpc}^{exc}$ | 1,2 | $1.25 \times 10^{-3}$ | Learning rate of STIM→HPC synapses |
| | 3,4,5,6,7 | $1.5 \times 10^{-3}$ | |
| $\lambda_{stim \rightarrow hpc}^{\beta}$ | 1,2 | 35 | $\beta_{stim \rightarrow hpc}/\eta_{stim \rightarrow hpc}^{exc}$ ratio for STIM→HPC synapses |
|  | 3,4,5,6,7 | 30 |  |
| $\tau_{stim \rightarrow hpc}^{cons}$ | All | 3 h | Synaptic consolidation time constant for STIM→HPC synapses |
| HPC→CTX |  |  |  |
| Parameter | Figure | Value | Description |
| $\eta_{hpc \rightarrow ctx}^{exc}$ | 2A-F | $1 \times 10^{-3}$ | Learning rate of HPC→CTX synapses |
| $\lambda_{hpc \rightarrow ctx}^{\beta}$ | 2A-F | 85 | $\beta_{hpc \rightarrow ctx}/\eta_{hpc \rightarrow ctx}^{exc}$ ratio for HPC→CTX synapses |
| $\tau_{hpc \rightarrow ctx}^{cons}$ | 2A-F | 20 min | Synaptic consolidation time constant for HPC→CTX synapses |

| THL EE |  |  |  |
| --- | --- | --- | --- |
| Parameter | Figure | Value | Description |
| $\eta_{thl}^{exc}$ | 3,4,5,6,7 | $1.5 \times 10^{-3}$ | Learning rate of EE synapses in thalamus |
| $\lambda_{thl}^{\beta}$ | 3,4,5,6,7 | 50 | $\beta_{thl}/\eta_{thl}^{exc}$ ratio for EE synapses in thalamus |
| $\tau_{thl}^{cons}$ | 3,4,5,6,7 | 20 min | Synaptic consolidation time constant for EE synapses in thalamus |
| HPC→THL |  |  |  |
| Parameter | Figure | Value | Description |
| $\eta_{hpc \rightarrow thl}^{exc}$ | 3,4,5,6,7 | $1.5 \times 10^{-3}$ | Learning rate of HPC→THL synapses |
| $\lambda_{hpc \rightarrow thl}^{\beta}$ | 3,4,5,6,7 | 50 | $\beta_{hpc \rightarrow thl}/\eta_{hpc \rightarrow thl}^{exc}$ ratio for HPC→THL synapses |
| $\tau_{hpc \rightarrow thl}^{cons}$ | 3,4,5,6,7 | 20 min | Synaptic consolidation time constant for HPC→THL synapses |
| THL→CTX |  |  |  |
| Parameter | Figure | Value | Description |
| $\eta_{thl \rightarrow ctx}^{exc}$ | 3,4,5,6,7 | $1 \times 10^{-3}$ | Learning rate of THL→CTX synapses |
| $\lambda_{thl \rightarrow ctx}^{\beta}$ | 3,4,5,6,7B | 105 | $\beta_{thl \rightarrow ctx}/\eta_{thl \rightarrow ctx}^{exc}$ ratio for THL→CTX synapses |
|  | 7C-F | 160 |  |
|  | 7G-J | 50 |  |
| $\tau_{thl \rightarrow ctx}^{cons}$ | 3,4,5,6,7 | 20 min | Synaptic consolidation time constant for THL→CTX synapses |
| All Excitatory Synapses with Long-Term Plasticity |  |  |  |
| Parameter | Figure | Value | Description |
| $\lambda^{\delta}$ | All | 0.02 | $\delta/\eta^{exc}$ ratio |
| $A$ | All | 1 | LTP rate |
| $\tau^{+}$ | All | 20 ms | Time constant of presynaptic trace for excitatory plasticity |
| $\tau^{-}$ | All | 20 ms | Time constant of postsynaptic trace for excitatory plasticity |
| $\tau^{slow}$ | All | 100 ms | Time constant of slow postsynaptic trace for excitatory plasticity |
| $P$ | All | 20 | Potential strength |
| $w^P$ | All | 0.5 | Upper fixed point of reference weight potential |
| $\tilde{w}$ | All | 0.0 | Initial reference weight |
| $\tau^{hom}$ | All | 10 min | Time constant of homeostatic regulation |
| $\tau^{ht}$ | All | 100 ms | Time constant of postsynaptic trace for homeostatic regulation |
| $w_{exc}^{max}$ | All | 5.0 | Maximum excitatory synaptic weight |
| $w_{exc}^{min}$ | All | 0.0 | Minimum excitatory synaptic weight |

| Inhibitory Synaptic Plasticity Model |  |  |  |
| --- | --- | --- | --- |
| Parameter | Figure | Value | Description |
| $\lambda^{\eta}$ | All | 50 | $\eta^{exc}/\eta^{inh}$ ratio |
| $\gamma$ | All | 4 Hz | Target activity level for cortex, hippocampus, and thalamus |
| $\tau^{iSTDP}$ | All | 20 ms | Time constant of pre- and postsynaptic traces for inhibitory plasticity |
| $\tau^H$ | All | 10 s | Time constant of global secreted factor |
| $w_{inh}^{max}$ | All | 5.0 | Maximum inhibitory synaptic weight |
| $w_{inh}^{min}$ | All | 0.0 | Minimum inhibitory synaptic weight |

| Stimulus Model |  |  |  |
| --- | --- | --- | --- |
| Parameter | Figure | Value | Description |
| $\nu^{bg}$ | All | 5 Hz | Background firing rate of stimulus population except in the consolidation phase |
| $\nu^{con}$ | 1,2 | 2 Hz | Background firing rate of stimulus population in the consolidation phase |
|  | 3,4,5,6,7 | 0 Hz |  |
| $\nu^{stim}$ | All | 25 Hz | Firing rate of neurons matching given stimulus when that stimulus is presented |
| $T_{On}^{training}$ | All | 1 s | Mean stimulus-on period in the training phase |
| $T_{Off}^{training}$ | All | 2 s | Mean stimulus-off period in the training phase |
| $T_{On}^{testing}$ | 1,2 | 1 s | Mean stimulus-on period in the testing phase |
|  | 3,4,5,6,7 | 2 s |  |
| $T_{Off}^{testing}$ | All | 2 s | Mean stimulus-off period in the testing phase |

| External Populations Model |  |  |  |
| --- | --- | --- | --- |
| Parameter | Figure | Value | Description |
| $\nu_{ext}^{bg}$ | 3,4,5,6,7 | 0 Hz | Firing rate of external populations except in the consolidation phase |
| $\nu_{ext}^{con}$ | 3,4,5,6,7 | 1 Hz | Firing rate of external populations in the consolidation phase |

| Network Simulation |  |  |  |
| --- | --- | --- | --- |
| Parameter | Figure | Value | Description |
| $T_{burn}$ | All | 120 s | Duration of burn-in period prior to training |
| $T_{training}$ | 1,2 | 45 min | Duration of training phase |
|  | 3,4,5,6,7 | 30 min |  |
| $T_{consolidation}$ | 1,2 | 12 h | Duration of consolidation phase |
|  | 3,4,5,6,7 | 24 h |  |
| $T_{testing}$ | All | 60 s | Duration of testing phase |
| $\Delta$ | All | 0.1 ms | Time step for updating neuronal state variables except for reference weights $\tilde{w}$ |
| $\Delta_{long}$ | All | 1.2 s | Time step for updating reference weights $\tilde{w}$ |

### Simulation of two-region networks with HPC and CTX

Simulations with two-region networks (i.e., Fig. 1**A**, Fig. **2A/G**) follow a defined sequence: burn-in, training, consolidation, and testing. The initial brief burn-in period of duration *T_burn_* stabilizes activity in each subnetwork under STIM background firing at rate *ν^bg^*. Subsequently, four random stimuli (depicted in Fig. 1**A**) are randomly presented to the network in the training phase of duration *T_training_* with equal probability and with inter-stimulus interval and stimulus presentation duration drawn from exponential distributions with means 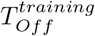 and 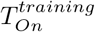, respectively. This is accomplished by maintaining the STIM background firing at *ν^bg^* but selectively increasing the firing rate of the STIM neurons that correspond to a given stimulus to *ν^stim^* for the duration of its presentation. Each stimulus consists of a non-overlapping random subset of 25% of the STIM neurons. Post-training, the network evolves spontaneously in the consolidation phase of duration *T_consolidation_* in the absence of stimulus presentations with STIM sustaining background firing at *ν^cons^.* It has been shown that reactivations of past experiences can take place during awake periods in both CTX and HPC [77, 78] and, hence, awake states may also be suitable for consolidation. However, our model aims capture recent findings that coupled oscillatory hippocampal-thalamic-cortical activity during sleep is essential for systems consolidation [30] and, hence, we set separate periods for training and consolidation. After consolidation, the network advances to the final testing phase of duration *T_testing_*. During testing, we present partial cues (depicted in Fig. 1**A**) to the network by keeping STIM background firing at *ν^bg^* and increasing the firing rate of the cue neurons to *ν^stim^*. Cue-off and cue-on periods also follow exponential distributions with means 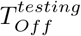 and 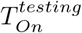, respectively. Each cue consists of a random 50% of the original stimulus. In the two-region networks, feedforward synapses have short- and long-term plasticity with the exception of HPC→CTX synapses in Fig.**2G** that only exhibit short-term plasticity. When blocking the output of engram cells in a given region, the inter-region efferent synapses of those cells are blocked but the recurrent counterparts are not to avoid finite-size effects. This procedure effectively allows for probing the effect of engram cells in downstream regions without changing the local engram dynamics. Critically, we set 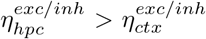 and 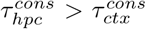. The higher learning rate in HPC relative to CTX reflects the experimental observation that engram cells in HPC are generated in an active state while those in CTX are initially in a silent state [33]. On the other hand, the longer synaptic consolidation timescale in HPC compared to CTX is intended to render newly-encoded engrams in HPC less stable than those in CTX consistent with reported engram dynamics [33]: engram cells in HPC switch from active to silent while those in CTX change from silent to active. In Fig. 2**A**, we set 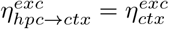 and 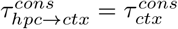. In Fig. 2**G**, HPC→CTX synapses are fixed (i.e., only have short-term plasticity). For a complete list of parameters for simulations of the two-region networks, see Table 1.

### Simulation of three-region network with HPC, THL, and CTX

Simulations with the three-region network (Fig. 3**A**) follow the same sequence as those with two-region networks (i.e., burn-in, training, consolidation, and testing). However, the three-region network is trained with four horizontal bars (as opposed to random stimuli) and is tested with partial cues consisting of the central 50% of the full bars (full bars and cues depicted in Fig. 3**A**). Furthermore, STIM does not provide background firing in the consolidation phase to reflect the gating of sensory processing by spindles during sleep [79–81]. Instead, random background input at rate 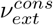 is provided independently to HPC and CTX during consolidation via two separate external populations of 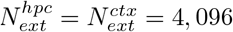 Poisson neurons. This reflects previous observations that THL activity is increased around hippocampal ripples coupled to spindles but suppressed otherwise [49] and that oscillations in HPC and CTX can occur independently of each other [50]. Outside consolidation periods, the external populations projecting to HPC and CTX remain silent (i.e., 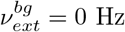). The procedure to block the output of engram cells in the three-region network is the same as in the two-region configuration. When blocking the output of inhibitory neurons, their efferent synapses onto both inhibitory and excitatory neurons are blocked. In simulations with HPC ablation (Fig. 7**A**), HPC and all its afferents and efferents synapses are removed from the network for the entirety of the testing phase. In the three-region network, feedforward synapses exhibit short- and long-term plasticity with the exception of those from STIM to THL (i.e., STIM→THL synapses only have short-term plasticity) as we assume that THL receptive fields have been learned during development and only change over timescales longer than those captured in our simulations. We set 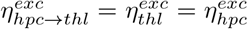 and 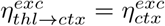 with 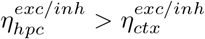. This is based on the view that the rate of change of synaptic weights tends to be higher for subcortical synapses compared to cortical ones. All plastic excitatory synapses share the same 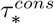 except the ones from STIM to HPC for which 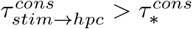. This is in line with the results of our simulations of two-region networks (Fig. 1) which showed that having a longer 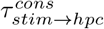 leads to a post-training decrease in feedforward STIM weights to HPC and the resulting de-maturation of its engram cells. For a complete list of parameters for simulations of the three-region network, see Table 1.

### Labelling engram cells and computing recall metrics

Engram cells are labelled in our model by computing the average stimulus-evoked firing rate of each neuron in a given subnetwork (i.e., HPC, CTX, or THL). A neuron is said to be an engram cell encoding a given stimulus if its average stimulus-evoked firing rate is above a threshold *ζ^thr^* = 10 Hz for the last Δ*t^eng^* = 300 seconds of the training phase. As a result, a single neuron may become an engram cell encoding multiple stimuli. In addition, an engram cell ensemble encoding a given stimulus is taken as activated upon presentation of a partial cue if its population firing rate is above the threshold *ζ^thr^* = 10 Hz during cue presentation. We then define recall true positive rate as the number of instances when the corresponding engram cell ensemble was activated following cue presentation divided by the total number of cue presentations in the testing phase. Inversely, recall false positive rate is computed by summing over all cue presentations the ratio of the number of incorrectly activated engram cell ensembles in response to a cue and the total number of non-matching engram cell ensembles (i.e., the number of stimuli minus one). Successful recall is said to happen when *only* the corresponding engram cell ensemble is activated by a partial cue (i.e., all other engram cell ensembles must be inactive). We then define recall accuracy as the number of successful recalls divided by the number of cue presentations in the testing phase. We compute 90% confidence intervals using a non-parametric bootstrap to aid in the visualization of the recall metrics.

### Simulation and data analysis details

We use the forward Euler method to update neuronal state variables with a time step Δ = 0.1 ms (except in the case of reference weights 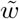 for which we use a longer time step Δ_*long*_ = 1.2 s for efficiency reasons). Population activity is computed with a temporal resolution of 10 ms without smoothing or convolution.

### Code

Code used to perform all simulations is written in C++ utilizing the Auryn framework for spiking neural network simulation [82]. Code used to analyze simulation results is written in Python. Simulation and data analysis code will be made publicly available upon publication.

## Acknowledgments

We thank Dheeraj S. Roy for feedback on an earlier version of this manuscript. This work was funded by the President’s PhD Scholarship from Imperial College London. This work was also supported by BBSRC BB/N013956/1, BB/N019008/1, Wellcome Trust 200790/Z/16/Z, Simons Foundation 564408, and EPSRC EP/R035806/1.

## Author contributions

D.F.T., S.S., and C.C. conceived the study and wrote the paper. D.F.T. performed the simulations and data analysis.

## Supplementary Material

**Figure S1.**
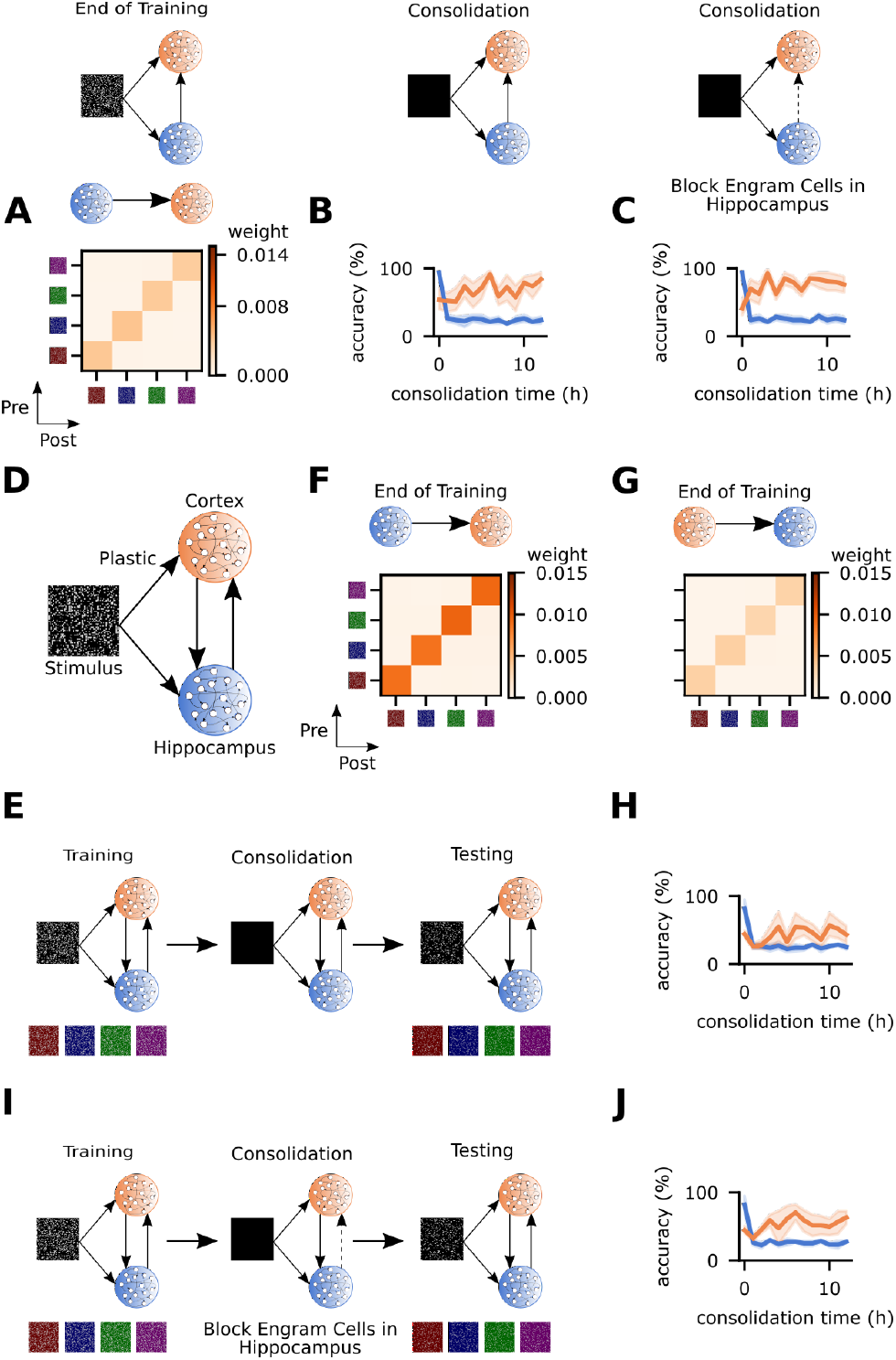
Monosynaptic HPC→CTX projections lead to inconsistent engram dynamics. Analysis of alternative network configurations with direct HPC→CTX synapses. **A-C**, Analysis of network in Fig. **2A** with 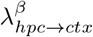 = 170. **A**, Mean HPC→CTX weight strength at the end of training clustered according to engram cell preference. **B-C**, Memory recall accuracy in the testing phase of the protocols in Fig. **2B** and **E**, respectively. **D**, Schematic of network model with plastic HPC→CTX and CTX→HPC synapses. Network and simulation parameters are the same as in Fig. **2A** except that 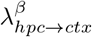 = 100 and *T_training_* = 35 min. Parameters for CTX→HPC are the same as HPC→CTX except *ϵ_ctx→hpc_* = 0.01. **E**, Schematic of simulation protocol with intact (control) HPC→CTX synapses for network **D**. **F-G**, Mean weight strength at the end of training clustered according to engram cell preference for network **D**. **F**, HPC→CTX. **G**, CTX→HPC. **H**, Memory recall accuracy in the testing phase of protocol **E**. **I**, Schematic of simulation protocol with the output of engram cells in HPC blocked during consolidation for network **D**. **J**, Memory recall accuracy in the testing phase of protocol **I**. **B-C**, Color as in Fig. 2**A**. **H/J**, Color as in **D**. **A/F-G/E/I**, Stimuli as in Fig. 1**A**. **B-C/H/J**, n = 5 trials with 90% confidence intervals.

**Figure S2.**
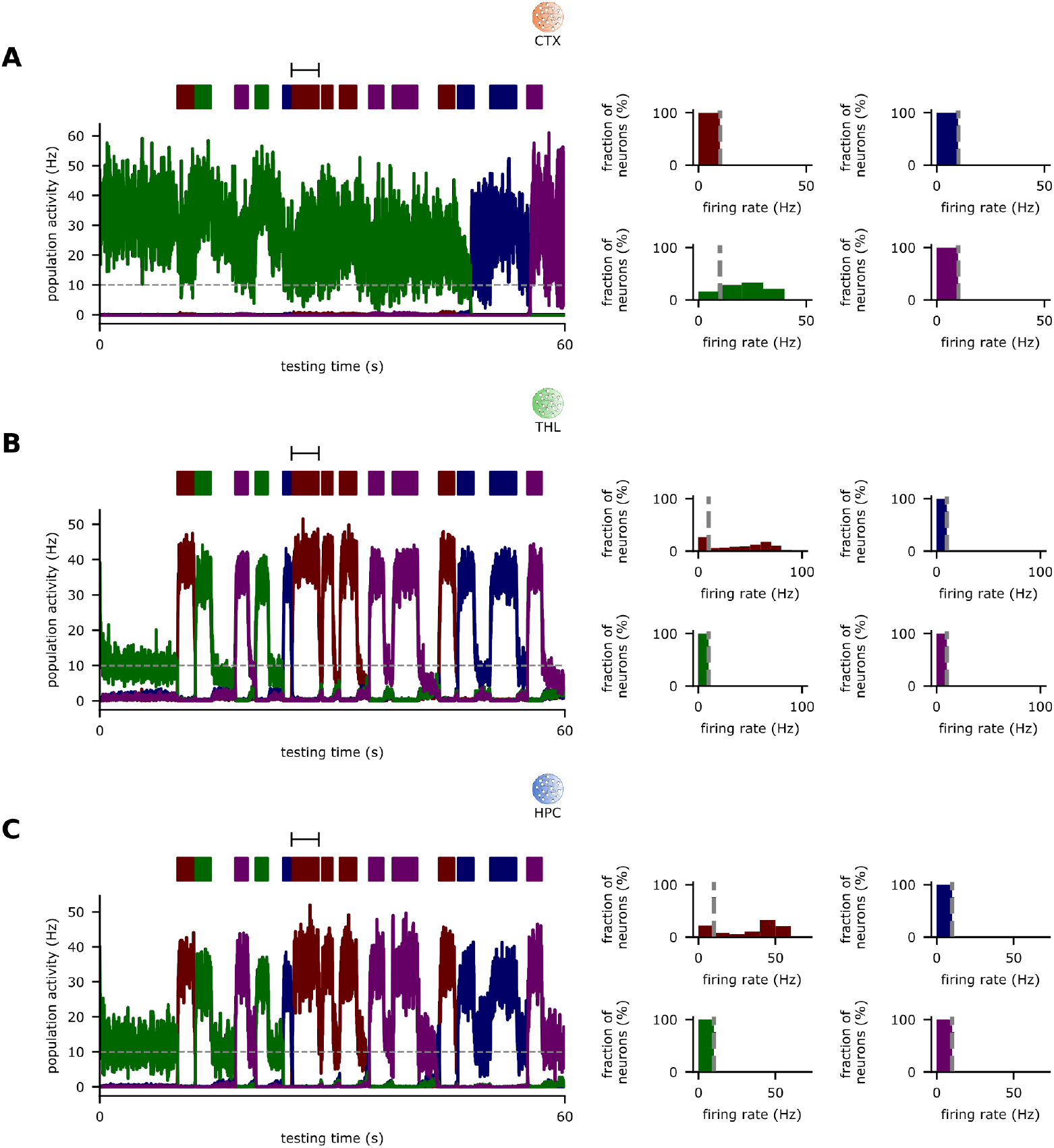
Recent memory recall relies on HPC engram cells. Analysis of recent memory recall in Fig. 3**B**. **A-C**, Left: Population activity of excitatory engram cells in the testing phase immediately following training (i.e., prior to consolidation) with cue presentation times displayed at the top. Right: histograms of the firing rates of engram cells encoding each stimulus for the cue presentation interval marked in the activity plot on the left. Dashed line in activity plots and histograms indicates threshold *ζ^thr^* = 10 Hz for engram cell activation. **A**, CTX. **B**, THL. **C**, HPC. **A-C**, Color as in Fig. 3**A**.

**Figure S3.**
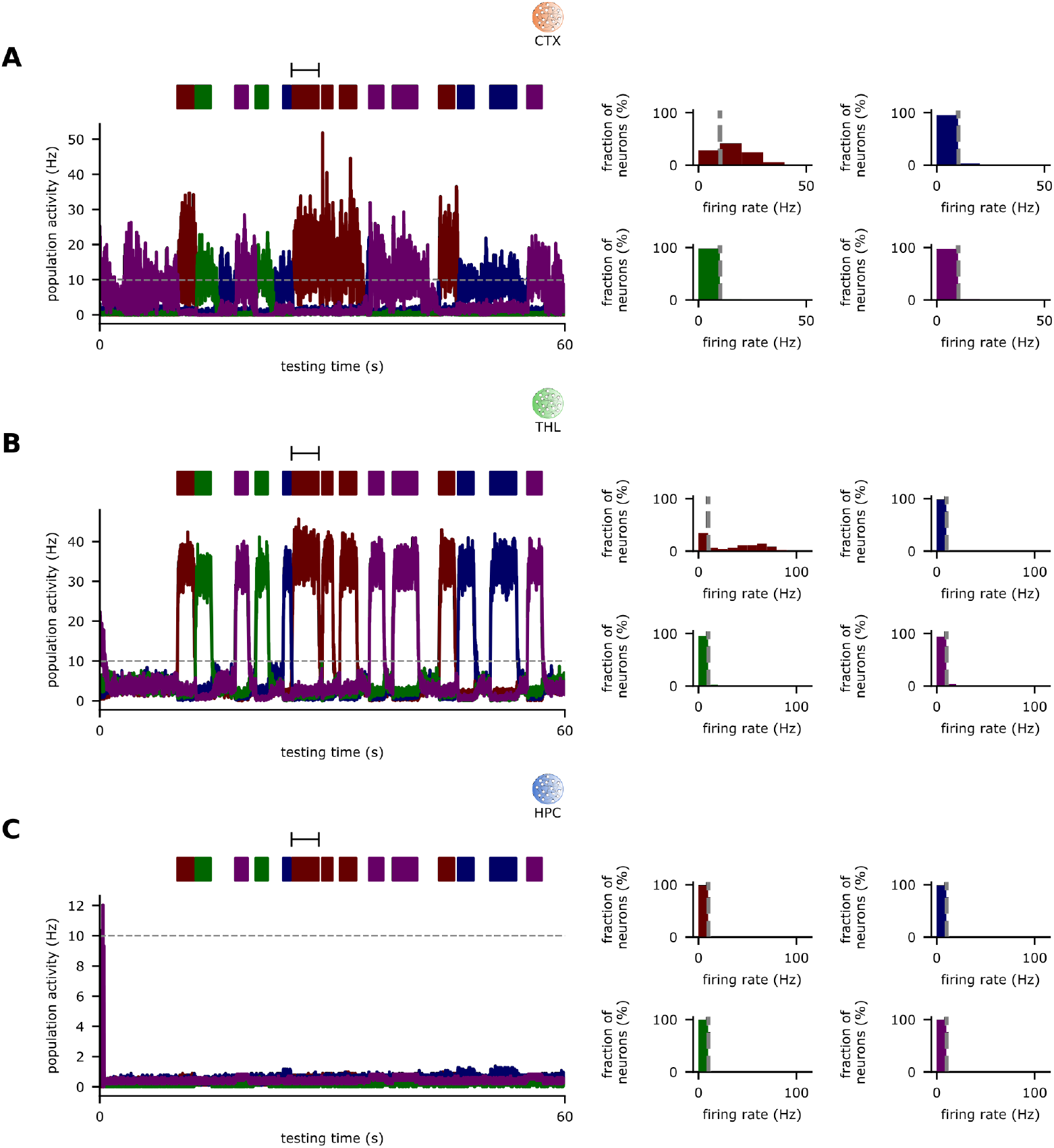
Remote memory recall relies on CTX engram cells. Analysis of remote memory recall in Fig. 3**B**. **A-C**, Left: Population activity of excitatory engram cells in the testing phase after 24 hours of consolidation with cue presentation times displayed at the top. Right: histograms of the firing rates of engram cells encoding each stimulus for the cue presentation interval marked in the activity plot on the left. Dashed line in activity plots and histograms indicates threshold *ζ^thr^* = 10 Hz for engram cell activation. **A**, CTX. **B**, THL. **C**, HPC. **A-C**, Color as in Fig. 3**A**.

**Figure S4.**
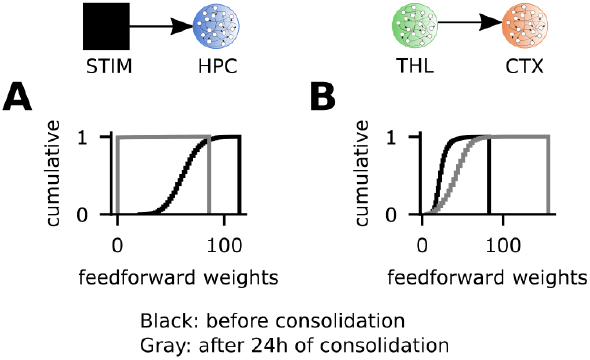
Feedforward weight changes underlie engram cell state transitions in HPC and CTX. Analysis of STIM→HPC and THL→CTX feedforward weights in Fig. 3**B**. **A-B**, Cumulative distribution function of the total feedforward synaptic weights onto individual excitatory engram cells at the end of training and after 24 hours of consolidation. **A**, Feedforward weights projecting from STIM to HPC excitatory engram cells. **B**, Feedforward weights projecting from THL to CTX excitatory engram cells.

**Figure S5.**
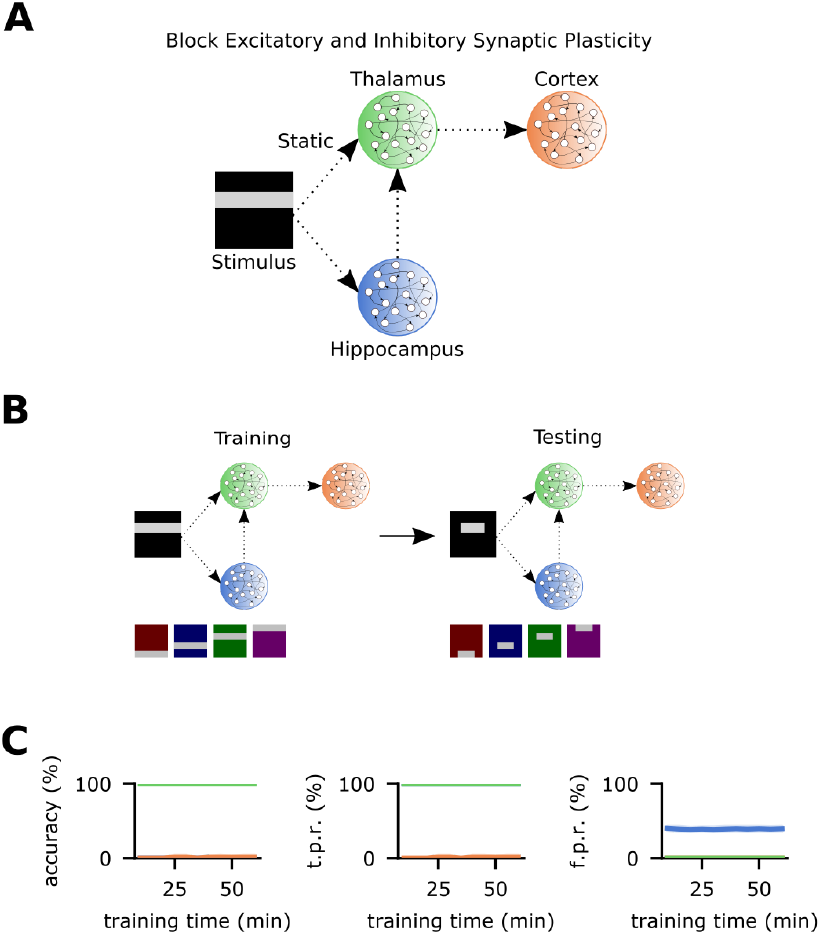
Excitatory and inhibitory synaptic plasticity are essential for memory acquisition. **A**, Schematic of network model with excitatory and inhibitory synaptic plasticity blocked in the entire network. **B**, Schematic of simulation protocol. Training and testing stimuli as in Fig. 3**A**. **C**, Memory recall in the testing phase as a function of training time. Recall curves (from left to right): accuracy, true positive rate, and false positive rate. Color as in **A**. **C**, *n* = 5 trials with 90% confidence intervals.

**Figure S6.**
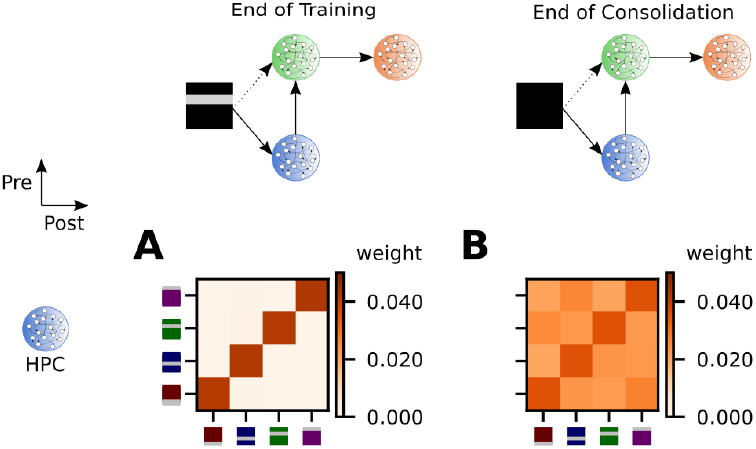
Silent HPC engram cells preserve recurrent excitatory connectivity. Analysis of HPC recurrent excitatory connectivity in Fig. 3**B**. **A-B**, Mean weight strength of recurrent excitatory synapses onto excitatory neurons in HPC clustered according to engram cell preference. **A**, At the end of the training phase. **B**, After 24 hours of consolidation.

**Figure S7.**
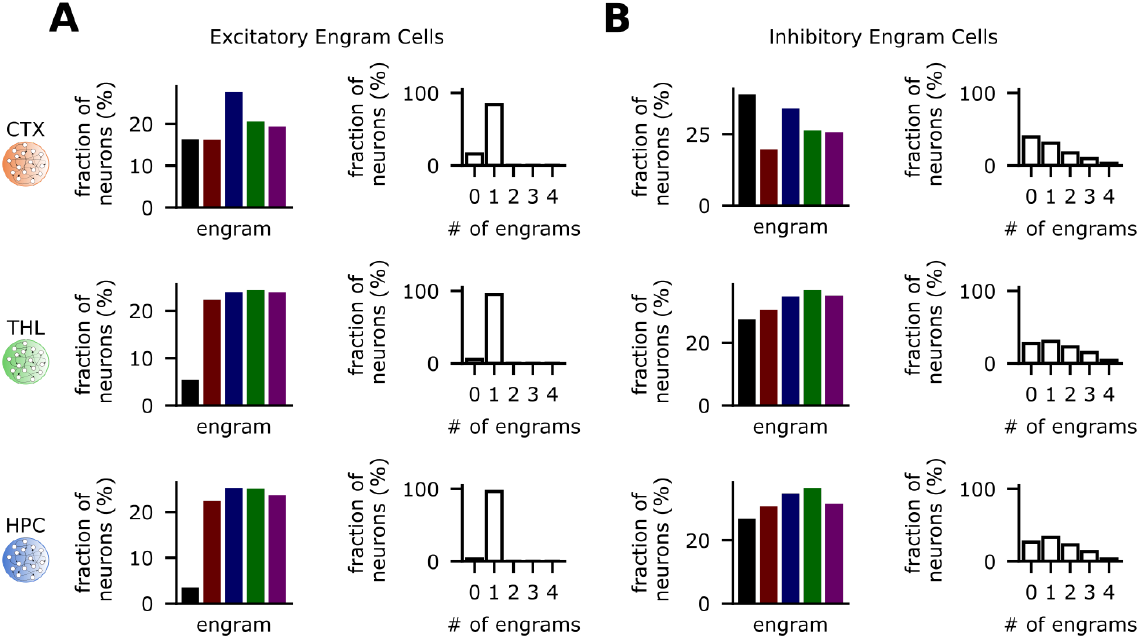
Composition of excitatory and inhibitory engram cells differ. Analysis of composition of excitatory and inhibitory engram cells in Fig. 3**B**. **A-B**, From left to right: proportion of neurons that encode each of the stimuli (black denotes no stimulus preference, other colors as in Fig. 3**A**), and proportion of neurons that encode 0-4 stimuli. From top to bottom: CTX, THL, and HPC. **A**, Excitatory engram cells. **B**, Inhibitory engram cells. Near-zero overlap in excitatory engram cells is a consequence of the zero overlap among training stimuli (Fig. 3**A**).

**Figure S8.**
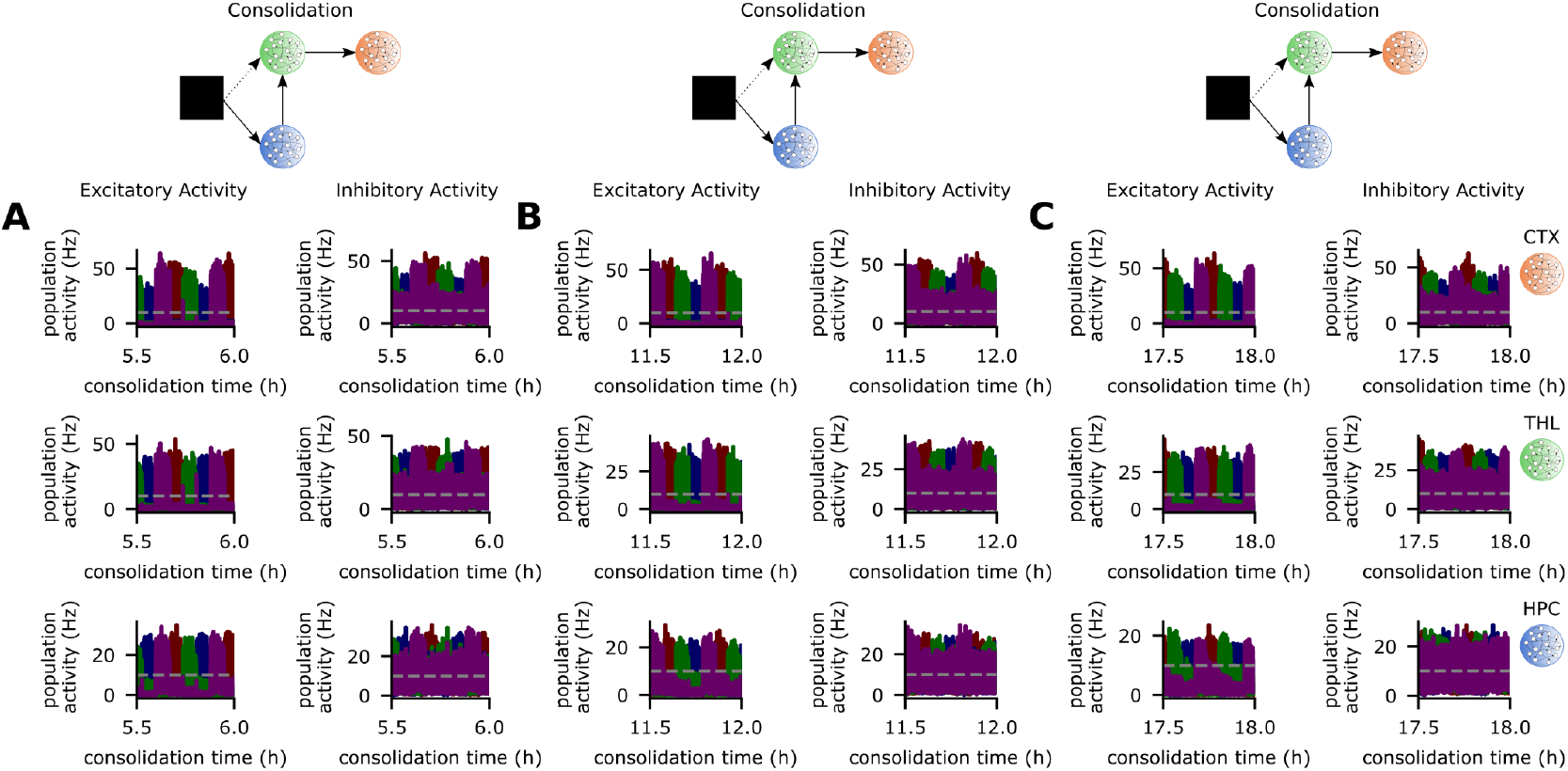
Coupled reactivations of excitatory and inhibitory engram cells throughout consolidation. Analysis of population activity of excitatory and inhibitory engram cells in Fig. 3**B**. **A-C**, Population activity of excitatory (left) and inhibitory (right) engram cells in the consolidation phase (dashed line indicates threshold *ζ^thr^* = 10 Hz for engram cell activation). Top to bottom: CTX, THL, and HPC. **A**, 5.5 to 6 hours of consolidation. **B**, 11.5 to 12 hours of consolidation. **C**, 17.5 to 18 hours of consolidation. **A-C**, Color as in Fig. 3**A**.

**Figure S9.**
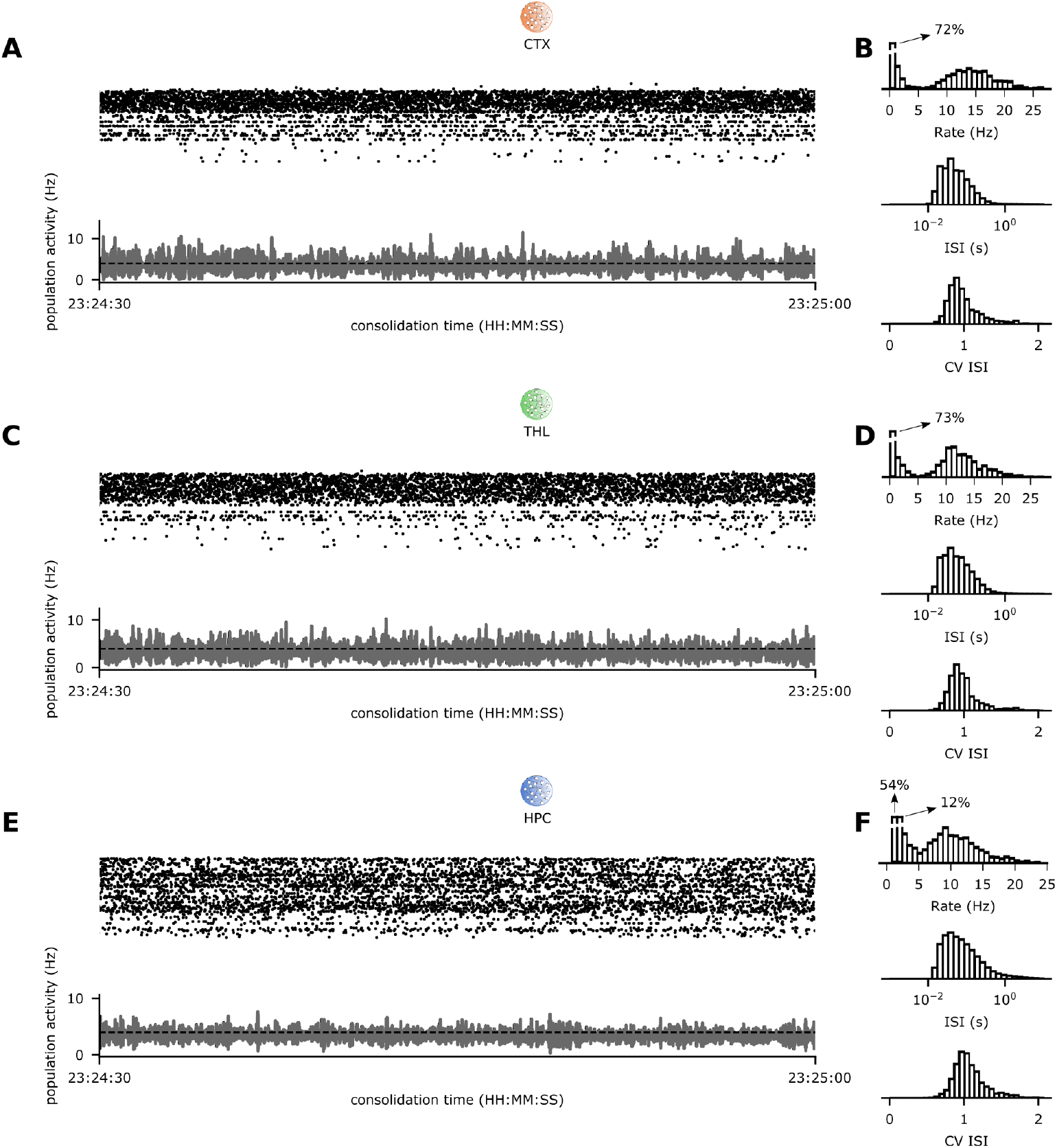
Inhibitory plasticity controls network activity in CTX, THL, and HPC. Analysis of excitatory network activity in Fig. 3**B**. **A/C/E**, Spike raster of random 256 excitatory neurons (top) and population activity of all excitatory neurons (bottom) in CTX (**A**), THL (**C**), and HPC (**E**) in a 30-second interval in the consolidation phase. For clarity, only every fifth spike is plotted in the raster. Dashed line in activity plots indicates target activity level γ = 4 Hz. Sample neurons with a higher firing rate in raster plots are part of engram reactivated in the time interval shown. **B/D/F**, Network statistics for **A**, **C**, and **E** respectively. From top to bottom: histograms of firing rates, interspike intervals, and coefficient of variation of interspike intervals (CV ISI).

**Figure S10.**
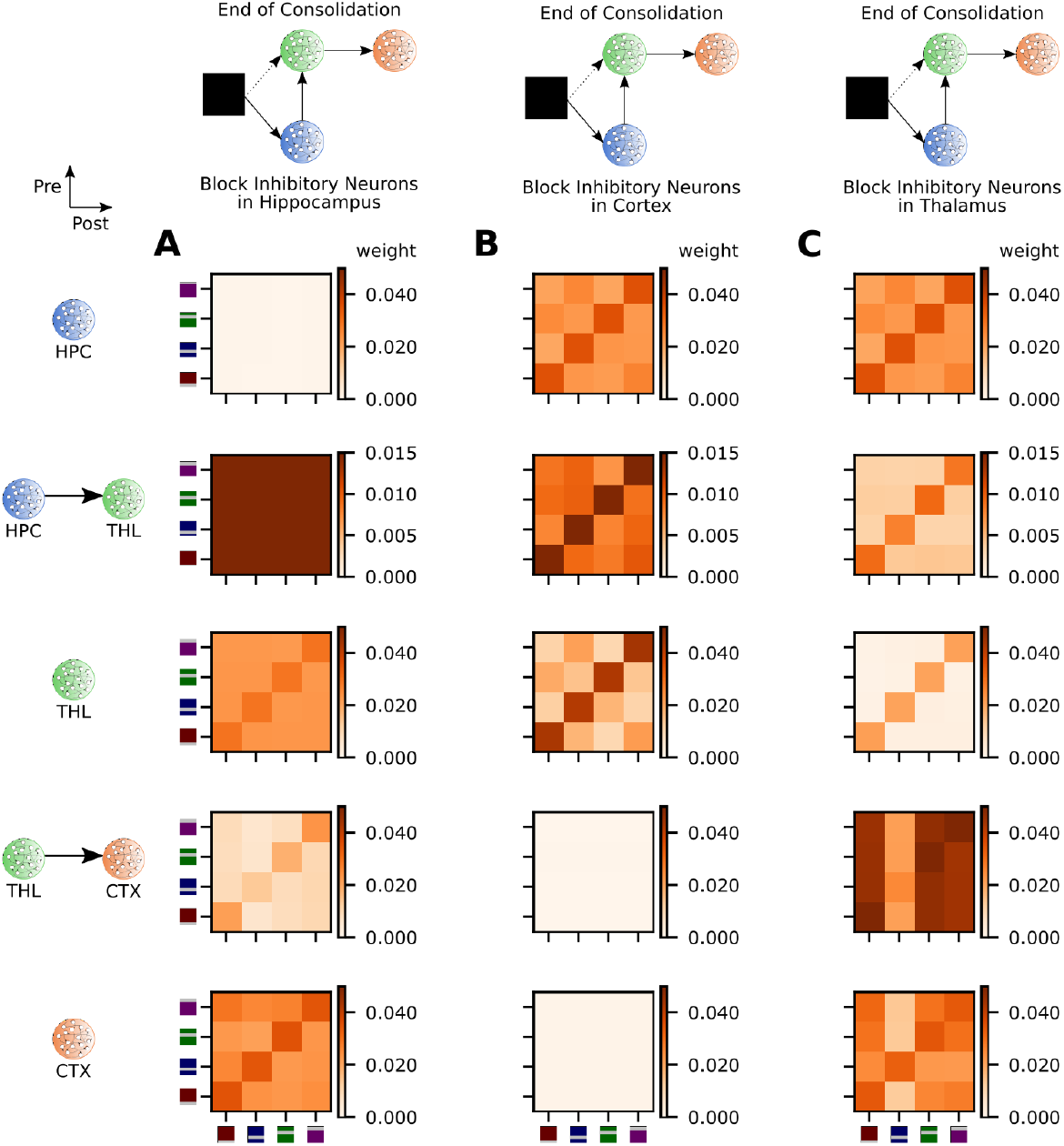
Inhibitory neurons are crucial for the consolidation of subcortical-cortical coupling. Analysis of synaptic coupling in Fig. 6. **A-C**, Mean weight strength of excitatory synapses onto excitatory neurons clustered according to engram cell preference. From top to bottom: HPC (recurrent), HPC→THL (feedforward), THL (recurrent), THL→CTX (feedforward), and CTX (recurrent). **A**, Mean weight matrices after 24 hours of consolidation for the network with blocked HPC inhibitory neurons (Fig. 6**A**). **B**, Mean weight matrices after 24 hours of consolidation for the network with blocked CTX inhibitory neurons (Fig. 6**B**). **C**, Mean weight matrices after 24 hours of consolidation for the network with blocked THL inhibitory neurons (Fig. 6**C**). **A-C**, Stimuli as in Fig. 3**A**.

